# A novel vaccine and drug targets for global eradication of bovine tuberculosis: Holistic frameworks for construction of a potent vaccine and identification of drug targets

**DOI:** 10.64898/2026.05.07.723640

**Authors:** Pooja Pawar, Sandhya Samarasinghe, Don Kulasiri

## Abstract

Bovine tuberculosis (TB), caused by *Mycobacterium bovis*, has become a global concern over the last two decades. Bovine TB primarily affects cattle, but other domestic livestock are also affected and it is more common in less developed and developing countries. The significant loss of livestock leads to trade restrictions and economic crises. Zoonotic potential of bovine TB raises health concerns for the public. Currently, no effective treatment is available and animal slaughtering is usually undertaken to reduce the burden of it in the environment. Antibiotic therapy can be used on animals living in captivity, but it is not reliable for herd or free-grazing animals. The BCG vaccine is another option available for treating the disease, but it shows limited efficacy in cattle. The prevention of bovine TB is a long-term goal that can only be accomplished by developing a more effective vaccine than BCG and designing new drugs. In this research, we propose therapeutic drug targets and vaccine for treating bovine TB. The conceptual framework for vaccine developed in this study uses a number of bioinformatics approaches to identify potential vaccine candidates and construct an in-silico epitope-based vaccine. Our holistic framework identified potential therapeutic candidates by directly analysing the proteome of TB bacterial strains. Specifically, we performed a comparative proteomic analysis of 11 *Mycobacterium bovis* strains to cover the diversity and identify conserved proteins among those strains for developing the bovine TB vaccine. An extensive reverse vaccinology and immunoinformatics analysis provided 26 highly immunogenic, non-toxic and non-allergenic epitopes (CTL epitopes-8, HTL epitopes-2 and B-cell epitopes-16) for *Mycobacterium bovis* required for three-dimensional structure construction of TB vaccine. The constructed epitope-based vaccine showed a potent interaction inside the host, thus generating efficient cell-mediated and humoral immune responses. Next, a framework based on a novel subtractive proteomic approach was developed for identifying bovine TB drug targets. We performed this approach on the 11 *Mycobacterium bovis* strains and identified nine drug targets that are conserved, essential, antigenic and have unique metabolic pathways in *Mycobacterium bovis*. These drug targets could further help investigate therapeutic drugs for the treatment of bovine TB. Several bioinformatics prediction tools were used together to ensure checks and balances, aiming to reduce the chance of errors and provide accurate results. The vaccine and drug targets developed in this study can be tested experimentally with confidence for further validation as therapeutics with the potential to eradicate bovine TB globally. The strategies implemented in the study are generic and can be used for other zoonotic infectious diseases. This study would be a game changer in the field of bovine tuberculosis treatment.

## 1. Introduction

Bovine tuberculosis is a chronic infectious disease that primarily affects cattle, but other livestock, such as deer, goats, horses and sheep, are also affected by the bacteria [1]. Bovine TB, a zoonotic disease, can spread to humans directly by inhaling aerosols or, indirectly, by ingesting unpasteurised milk. Affected global livestock population has reached 20-30%, resulting in economic losses of more than USD 3 billion annually across the world [1]. Bovine TB also decreases milk production by four and 20 per cent worldwide. There is a significant loss of livestock and trade restrictions due to bovine TB, e.g., in New Zealand, the beef and dairy industries are at potential risk of TB [2]. No effective treatment is available for bovine TB due to its infectious nature and the drug resistance of *Mycobacterium bovis*. Antibiotic therapy and vaccine (BCG) have shown limited effectiveness in cattle. The prevention of bovine TB is a long-term goal that can only be accomplished by developing a more effective vaccine than BCG and designing new drugs. Developing a bovine TB epitope-based vaccine containing B-cells and T-cells (MHC class I and MHC class II-restricted) epitopes could elicit a humoral and cell-mediated immune response. Thus, protective immunity is required to lower the transmission of infection. This study implemented comparative proteomic analysis, reverse vaccinology, immunoinformatics and structural vaccinology to design and develop an effective epitope-based bovine TB vaccine. A conceptual method to identify drug targets against *Mycobacterium bovis* was also employed in this study to answer pathogenicity and drug resistance against bovine TB. These drug targets could help treat bovine TB by using or modifying the available veterinary drugs.

Bovine tuberculosis is caused by *Mycobacterium bovis.* It is a rod-shaped, intracellular, aerobic, gram-positive and slow-growing bacterium. *Mycobacterium bovis* cell wall comprises covalently linked peptidoglycans, arabinogalactans, non-peptidoglycan amino acids and a glucan [3]. The cell structure and metabolism of *Mycobacterium bovis* is similar to *Mycobacterium tuberculosis*. The main *in-vitro* difference in *Mycobacterium bovis* is a point mutation in *pykA* that affects the binding of the Mg^2+^ cofactor with pyruvate kinase in the final step of glycolysis. The genome of *Mycobacterium bovis* is 99.95% identical to *Mycobacterium tuberculosis* [4]. A fully virulent strain, *Mycobacterium bovis* AF2122/97, was isolated from an infected cow in 1997 [4]. The complete information on the genome sequence of *Mycobacterium bovis* AF2122/97 was published in 2003. The *Mycobacterium bovis* genome contains a single, circular chromosome containing 4,345,492 base pairs with a high G+C content of 65.6% [4]. Around 4043 genes encode 3988 proteins in *Mycobacterium bovis* AF2122/97. Developments in high-throughput experimental methods have made completed genomes available for different *Mycobacterium bovis* strains.

Cattle are the primary host of infection, but *Mycobacterium bovis* can infect many hosts, including humans as well as domestic and wild mammals [1]. The reported susceptible domestic species are sheep, goats, pigs, deer, horses, camels, cats, dogs and ferrets (The Centre for Food Security and Public Health, Iowa State University). Wild mammals infected by *Mycobacterium bovis* are bears, African buffalo, elephants, raccoons, primates, possums, rhinoceros, foxes and rodents [5]. A maintenance host can be defined as a species endemic to infection that transmits the disease to other animals by direct contact [6]. There are several maintenance hosts for *Mycobacterium bovis,* such as brush-tail possums (New Zealand), badgers (the United Kingdom and Ireland), and bison and elk (Canada) [5]. *Mycobacterium bovis* is usually transmitted by the inhalation or ingestion of droplet nuclei. The infection can also be transmitted through bodily fluids such as urine, saliva, milk and colostrum. The transmission of *Mycobacterium bovis* depends on several factors such as the frequency of excretion, infective dose through coughing, period of communication with the infected host and the host’s susceptibility [7]. Humans get infected by *Mycobacterium bovis* by ingesting unpasteurised milk and raw or undercooked meat [1]. A person working in dairy farms or slaughterhouses is more susceptible to TB. The infected person can transmit the infection to other people, but the transmission rate of bovine in humans is low [8].

Bovine TB is found globally, but eradication programmes have helped eliminate or nearly eliminate this chronic disease from domestic animals in several countries. The countries reported as TB-free include Iceland, Denmark, Sweden, Switzerland, Norway, Canada, Singapore, Australia and Finland (The Centre for Food Security and Public Health, Iowa State University). The World Organization for Animal Health (OIE) has 181 member countries. In 2017, 78 countries reported the prevalence of bovine TB (World Organization for Animal Health, 2018). Reis et al. (2020) studies showed the prevalence of bovine TB is more in Ireland (22.87%) and the United Kingdom (16.43%) compared to France (5.03%) and Spain (3.05%) [9]. The disease prevalence is still high in Africa and some parts of Asia. Srinivasan et al., 2018 estimated that 21.8 million cattle are infected with bovine TB in India [10]. Currently, several eradication programmes focussing on eliminating the maintenance hosts are being implemented in the United Kingdom, the U.S.A, Mexico, Japan and New Zealand. The precise estimation of the prevalence of bovine TB remains poorly understood at the global level due to poor diagnosis of livestock and domestic animals and the lack of surveillance data from most countries in the world. *Mycobacterium bovis* infections detected in wildlife can cause severe health implications to other organisms living in the same ecosystem. Bovine tuberculosis has zoonotic potential, raising health concerns for the public [11]. Poor sanitation and no isolation of infected cattle from the herd on the farm can lead to exposure of *Mycobacterium bovis* to humans. The pasteurization of milk is not compulsory in India. Thus, bovine TB in cattle also impacts human health. It is also important to remember that getting infected from eating the meat of the infected animal is less likely, but the risk is still there. Out of 10 million incidence cases in humans in 2019, WHO estimated 0.14 million cases were zoonotic TB caused by *Mycobacterium bovis*, with 11,400 human deaths [12].

The immune response generated by cattle against *Mycobacterium bovis* is similar to the human immune response against *Mycobacterium tuberculosis* because of the similarity of the mammalian immune system and cells. In cattle, a primary infection results in a lesion in the nasopharynx and upper parts of the lungs [13]. The alveolar macrophage is the primary host for intracellular growth of *Mycobacterium bovis* and performs phagocytosis. In phagocytosis, the alveolar macrophage ingests *Mycobacterium bovis* forming a phagosome, which later forms a phagolysosome complex together with a lysosome. The formation of phagolysosome causes the breakdown of *Mycobacterium bovis* into smaller fragments with the help of hydrolytic enzymes present in the lysosome. In the lung, alveolar macrophages play a vital role in interacting with innate and acquired immune cells. After initial infection, the alveolar macrophages present the bacterial fragments (antigens) to T-helper cells leading to activation of cell-mediated immunity. The release of anti-inflammatory cytokines IL-4, IL-5, IL-10 and IL-13 by T-helper cells promotes the activation of B-lymphocytes, leading to antibody production. The B-cells and T-cells surround the site of primary infection and form a mass of tissue called a granuloma. In the initial stages of bovine TB infection, cattle are asymptomatic, so it can take a few months to years for any sign or symptom to be observed. As the infection progresses, the infected animal shows clinical signs that include weakness, fluctuating fevers, prominent lymph nodes, a moist cough with increased breathing rate, loss of weight, emaciation, diarrhoea, and induration of the udder.

The techniques involved in controlling bovine tuberculosis in wildlife are restricted. The isolation of infected animals is not an option in hugely populated or low-income countries. In New Zealand, possums serve as a significant wildlife reservoir for *Mycobacterium bovis* infection, and, since 1994, there has been a marked increase in the implementation of possum poisons [14]. This resulted in a 70% reduction in *Mycobacterium bovis*-infected cattle herds from June 1994 to June 2001. Animal test-and-slaughter schemes implemented in several countries have reduced the prevalence of bovine tuberculosis. Still, such expensive control programmes have raised economic burdens and increased opposition by the farmers [15,16].

Currently, there is no effective treatment available for bovine TB due to its infectious nature and drug resistance of *Mycobacterium bovis*. The available treatment of bovine TB mainly depends on the health status of the infected species. Antibiotic therapy can be used for animal species living in captivity, but this is not reliable for herd or free-grazing animals. *Mycobacterium bovis* is naturally resistant to pyrazinamidase (first-line TB drug) [17]. First-line human TB drugs for treating livestock are also ineffective and costly as the treatment requires six to nine months of daily medication. BCG vaccine is another option for treating the disease, but it shows limited efficacy in cattle. BCG is prepared from a live-attenuated strain of *Mycobacterium bovis* [18]. It is produced using living *Mycobacterium bovis*, which has been weakened or attenuated under specific laboratory conditions. While it cannot cause the disease, it retains the ability to generate an immune response within the host. BCG has been around for almost a hundred years and was first used in 1921 [19], yet BCG vaccine has not provided a sufficient level of protection. Several attempts have been made to develop a live-attenuated or heat-killed vaccine against bovine TB in cattle, but none has been successful [20,21,22,23].

As the lives of humans and animals are interrelated, it is essential to implement specific therapeutic measures to reduce the burden of bovine TB globally. This study aims to identify potential vaccine and drug targets for bovine TB treatment using several bioinformatics approaches. The current explosion in bioinformatics has revolutionized the field of vaccine and drug development. It provides new tools that facilitate the identification of potential vaccine and drug targets without the need to culture the pathogens in the laboratory. Information about the genome, transcriptome or proteome of *Mycobacterium bovis* can help identify novel therapeutic candidates.

### 1.1 Aims

Vaccination is still a promising strategy to protect livestock and reduce the incidence rate of bovine TB. Vaccinating cattle against tuberculosis aims to prevent infection in these animals. Interest in developing and using TB vaccines for cattle can be rekindled with an improved understanding of the protective immunity against TB. The development of a new vaccine against bovine TB is considered a control option to address two important issues: preventing the establishment of the disease and eliminating zoonotic bovine TB. This study aims to design an improved vaccine that can protect against bovine TB disease. Generating effective memory cells against bovine TB would be a crucial step toward tackling the disease worldwide. Epitope-based vaccine is a powerful strategy for this. The epitope-based bovine TB vaccine containing B-cell and T-cell (MHC class I and MHC class II-restricted) epitopes could elicit a humoral and cell-mediated immune response. Compared to the conventional approach, comparative proteomic analysis, reverse vaccinology, immunoinformatics, and structural vaccinology (Figure 1) helps to develop a safer vaccine as a careful selection of non-toxic, non-allergenic immunodominant epitopes is undertaken. The main advantage of an epitope-based vaccine is the removal of deleterious epitopes that can cause cross-reactive reactions or autoimmunity in the host. Epitope-based vaccines are more potent, targeting multiple-conserved epitopes and, when controlled correctly, induce a specific immune response to a broad range of immunodominant epitopes and break immune tolerance.

**Figure 1:**
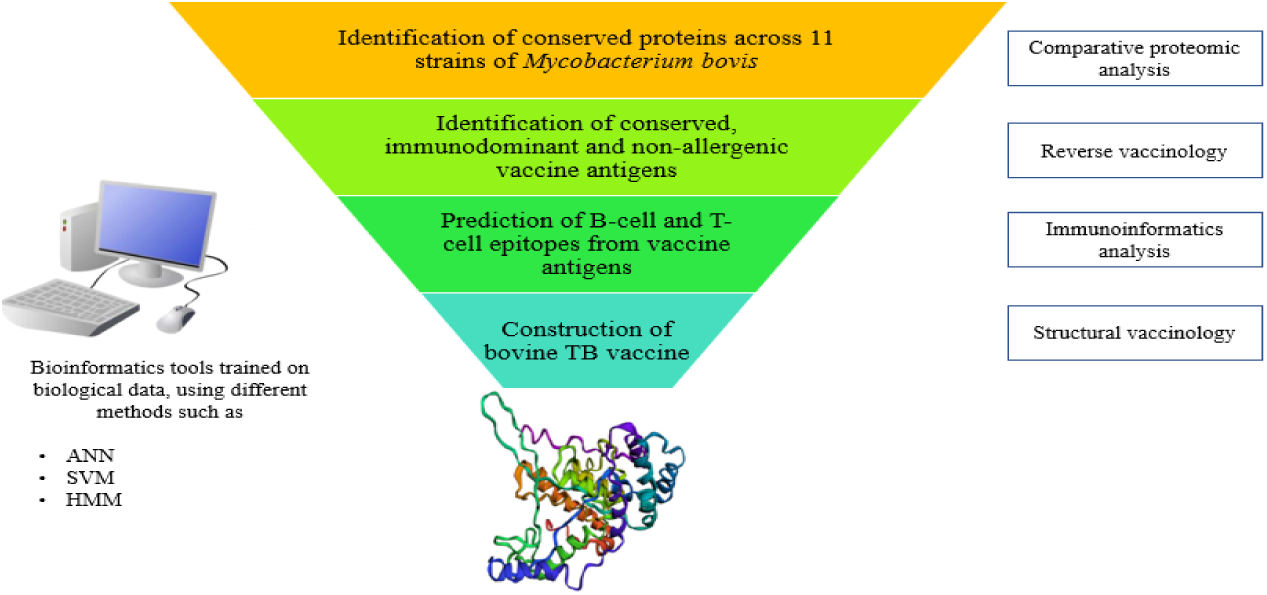
Conceptual framework for developing *in-silico* a vaccine against *Mycobacterium bovis*

To identify the therapeutic drug candidates for bovine TB treatment, a new bioinformatics approach was developed to answer pathogenicity and drug resistance against bovine TB. The emergence of antimicrobial resistance (AMR) has reduced the efficacy of antibiotics in treating the disease. An approach that identifies conserved and pathogenic drug targets to design better drug therapeutics against bovine TB was employed to overcome the antimicrobial issue. Various bioinformatics approaches can be used for examining the proteome of *Mycobacterium bovis* to identify potential drug targets that can facilitate drug development for bovine TB treatment. A subtractive proteomic approach was used to determine the conserved, essential, antigenic and druggable targets having unique metabolic pathways in *Mycobacterium bovis* (Figure 2).

**Figure 2:**
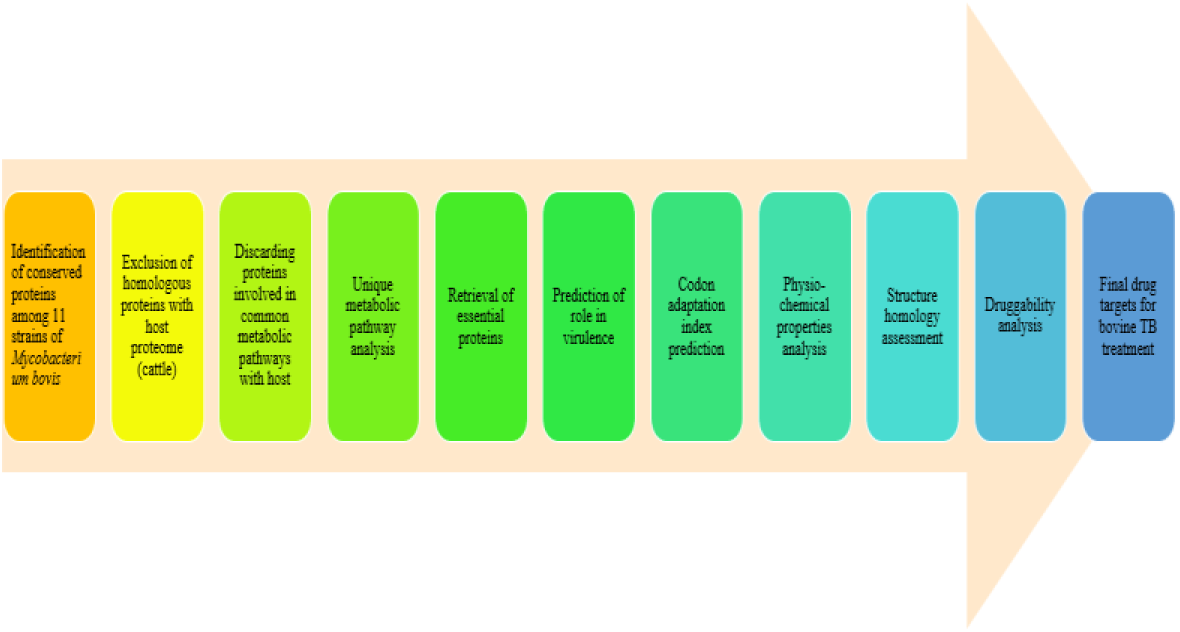
Comprehensive subtractive proteomic analysis for the identification of potential drug targets in *Mycobacterium bovis*

## 2. Materials and Methods

This section describes the step-by-step approach for designing a bovine TB vaccine and identifying potential vaccine candidates against bovine tuberculosis. Various bioinformatics tools were used for analysing the proteome of 11 *Mycobacterium bovis* strains.

### 2.1 Comparative proteomic analysis

The complete proteome sequence of 11 strains of *Mycobacterium bovis* was downloaded in FASTA format via the National Center for Biotechnology Information (NCBI) Genome FTP site. Blank spaces and unwanted information were removed, and multi-line protein sequences were converted into single-line FASTA format. Standalone BLAST (Basic Local Alignment Search Tool) [24,25] was downloaded from the NCBI FTP site for performing a homology search. *Mycobacterium bovis* AF2122/97 was used as a reference proteome to identify conserved proteins with more than 99% sequence similarity across all 11 strains. The conserved protein sequences identified were selected for further analysis for identifying potential therapeutic bovine TB vaccine and drug candidates to solve the problem of drug resistance and provide broad coverage.

### 2.2 Framework for epitope-based bovine TB vaccine design and development

#### 2.2.1 Subtractive reverse vaccinology analysis

The conserved proteins (results of previous steps) were examined using reverse vaccinology to identify outer-membrane, highly antigenic and non-allergenic proteins. These antigenic proteins would help design and develop an epitope-based bovine TB vaccine (Figure 1).

The proteins in the outer-membrane and extracellular space are considered suitable vaccine targets as they are easily accessible to the host immune cells. Using six different subcellular localisation (SCL) prediction tools (PSORTb v.3.0 [26], CELLO [27], LocTree3 [28], SOSUI [29], pLoc_bal-mGpos [30] and GramLocEN [31]), the proteins present in the cell membrane, cell wall and extracellular space were determined. The selected proteins were analysed for the presence of a transmembrane α-helix. Protein with more than one transmembrane α-helix is suitable for vaccine and drug development studies as they are easy to clone in the laboratory. TMHMM server predicts transmembrane regions and their orientation using a hidden Markov model (HMM) [32]. Proteins are exported across the cytoplasmic membrane with the help of secretory proteins. SignalP 4.1 [33] was used to identify proteins with signal peptides, and SecretomeP [34] was used to predict secretory proteins. PRED-TAT [35] was used for identifying TAT-signal peptides. Lipoproteins are highly immunogenic and antigenic targets in the *Mycobacterium tuberculosis* Complex (MTBC). PRED-LIPO was used for predicting potential lipoproteins [35].

The antigenic proteins are potential vaccine targets as they can stimulate a precise immune response in the host. VaxiJen [36], VirulentPred [37] and MP3 [38] tools were used to determine the antigenicity of the selected proteins. The proteins predicted antigenic by all three prediction tools were further used for allergenicity analysis. Algpred [39] uses several approaches (BLAST + IgE epitopes + MAST (Motif alignment and search tool) + Machine learning techniques + Ensemble approach) to determine the allergenic nature of the selected antigenic proteins. Non-allergenic proteins were selected. The adhesin proteins of mycobacteria help in its attachment to host cell-surface receptors. The adhesion probability was predicted by software for predicting adhesins and adhesin-like proteins using neural networks (SPAAN) [40]. The proteins with an adhesion probability score of 0.5 or above were chosen for further analysis.

The homologous proteins can cause hypersensitive reactions in the host body. For predicting homology between cattle and *Mycobacterium bovis*, BLASTp was used for identifying homologous proteins. Proteins with a sequence identity of 30% or more and a bit score of more than 100 were eliminated.

#### 2.2.2 Immunoinformatics analysis for prediction of B-cell and T-cell epitopes

The outer-membrane bovine proteins predicted as antigenic and non-allergenic were further analysed for predicting potential B-cell and T-cell epitopes using immunoinformatics tools to develop a potent vaccine eliciting a robust immune system response against bovine tuberculosis.

##### 2.2.2.1 B-cell epitope prediction

B-cell epitopes are necessary for instigating the humoral immune response. ABCpred was used for the prediction of the B-cell epitopes. ABCpred predicts epitopes with an accuracy of 65.93% using a recurrent neural network (RNN) [41]. The length of B-cell epitopes was set to 20 amino acid residues, with a threshold score of 0.5.

##### 2.2.2.2 T-cell epitope prediction

MHC molecules are present on the surface of antigen-presenting cells, such as macrophages and dendritic cells. The function of the MHC molecules is to present the fragmented or processed antigen to the appropriate T-cell (HTL or CTL) of the immune system. This initiates the cell-mediated immune response and secretion of the cytokines to eliminate the infection from the host.

###### (i) Prediction of MHC class I binding T-cell (cytotoxic-T cell) epitopes

MHC class I molecules present the epitope to cytotoxic T-lymphocytes (CTL). Selecting epitopes that bind strongly to MHC class I is the most crucial step in the vaccine design. It helps predict the most antigenic and immunodominant epitopes required to initiate the immune response. The following bovine MHC class I HLA alleles were used to predict MHC class I restricted T-cell epitopes: BoLA-D18.4, BoLA-AW10, BoLA-JSP.1, BoLA-HD6, BoLA-T2a, BoLA-T2b and BoLA-T2c. Two methods were employed in this step: IEDB MHC-I [42] and NetMHCpan 4.1 server [43]. In the IEDB MHC-I method, the prediction was made using an artificial neural network for determining the binding affinity of epitopes with bovine HLA alleles. Nine-mer epitopes with a percentile rank lower than 1.0 were chosen. The lower value of percentile rank indicates a higher affinity of CTL epitopes towards the MHC class I molecules. The NetMHCpan 4.1 server predicts the peptides binding to the MHC class I molecule. The method uses an artificial neural network trained with 201 different MHC alleles of humans, mice, cattle, primates and swine. The technique has shown a strong preference for the 9-mer peptide [43]. The epitopes commonly predicted by both tools were selected as T-cell MHC class I epitopes or CTL epitopes.

###### (ii) MHC class II binding T-cell (helper-T cell) epitope prediction

MHC class II molecules present epitopes to the naïve T-helper lymphocytes (HTL). This leads to the release of cytokines that facilitate the transformation of the naïve-T-helper cells into effector T-cells or memory T-cells. The method used for the prediction of HTL epitopes was the IEDB MHC II server. Using the consensus prediction method, IEDB MHC-II helped predict 15-mer MHC-II epitopes [44]. The HLA alleles of the mouse model were used in this step as no information was available for bovine MHC-II molecules. The epitopes with a percentile rank lower than 1.0 were chosen.

#### 2.2.3 Filtering of epitopes

B-cell and T-cell epitopes were then filtered to identify immunodominant antigenic, non-toxic and non-allergenic epitopes. VaxiJen [36] was used to predict the epitope’s potential to initiate an immune response based on the physiochemical properties of the query sequence. Epitopes with a VaxiJen score of 0.7 or above were selected. The toxic epitopes are more reactogenic inside host. Thus, prediction of non-toxic epitopes was done using the SVM-based prediction tool, ToxinPred [45]. Allergenicity prediction is crucial for avoiding allergens in designing the bovine TB vaccine. AllerTOP 2.0 predicts epitope allergenicity based on the input sequence’s physiochemical properties [46]. The allergenic epitopes were excluded from the study. The hydrophilic epitopes are present on the surface of the antigenic proteins. The selection of hydrophilic epitopes would help construct a vaccine that initiates a quick immune response. The grand average of hydropathicity (GRAVY) [47] was predicted using ProtParam.

#### 2.2.4 Bovine TB vaccine construction using structural vaccinology

After choosing the immunodominant B-cell and T-cell epitopes, structural vaccinology was employed to design a three-dimensional (3D) structural model for the bovine TB vaccine and evaluate its interaction with the toll-like receptors in the host (cattle).

##### 2.2.4.1 Vaccine design

To design a bovine TB vaccine sequence from the shortlisted epitopes, HTL, CTL and B-cell epitopes were joined with the help of a flexible linker, AAY. An adjuvant 50s ribosomal protein L7/L12 was attached using an EAAAK linker at the N-terminal of the bovine TB vaccine protein sequence.

##### 2.2.4.2 Analysis of antigenicity and physiochemical properties

VaxiJen [36] was used to predict the antigenicity of the final vaccine construct. Allergenicity prediction was made to exclude allergens. Algpred, an SVM model, was used to determine the vaccine protein’s allergenicity [39]. Different physiochemical properties of the final bovine TB vaccine were computed using Protparam. These properties include molecular weight (MW), isoelectric point (pI), amino acid composition, extinction coefficient [48], instability index (II), aliphatic index [49] and the grand average of hydropathicity (GRAVY) [47].

##### 2.2.4.3 Vaccine structural model construction

The secondary structure (alpha-helices, beta-sheet and coil) of the bovine TB vaccine construct was evaluated with the help of PSIPRED [50]. Interpreting secondary structure enhances the understanding of the vaccine protein’s tertiary structure model. The tertiary structure of the bovine TB vaccine was constructed using RaptorX [51]. RaptorX performs a template search based on the similarity of the input protein sequence and then constructs a good-quality structural model for the input query protein. The output structural model was refined using GalaxyRefine to reduce any distortions present in the structure of the vaccine protein [52]. The output contains five structural models, GDT-HA (global distance test-high accuracy), MolProbidity, clash score, RMSD, poor rotamers, and Rama favoured scores. PROCHECK and Verify 3D were used for structure validation. PROCHECK checks the stereochemical quality of the modelled protein [53]. Verify 3D evaluates the compatibility of the 3D atomic model with its amino acid sequences by assigning a structural class based on location and environment (α-helices, β-sheets and loops) [54]. If the vaccine structural model had more than 90% residues in the most favoured region, it was considered the best quality model and was used for further analysis.

##### 2.2.4.4 Vaccine-TLR molecular docking

Bovine toll-like receptors (TLRs) are present on the surface of antigen-presenting cells (macrophages or dendritic cells). TLRs interact with the pathogen/vaccine antigen to initiate the innate immune response. The structures of bovine TLRs were not present in the PDB database. The protein sequence of bovine used for structure construction was Q95LA9, Q9GL65 and B5T278 for TLR-2, TLR-4 and TLR-6, respectively. I-TASSER server was used for generating the 3D structure of TLRs. I-TASSER is an online platform for automated protein structure predictions from amino acid sequences. The I-TASSER suite pipeline consists of four essential steps [55]

- *Threading template prediction*: after submission of the query sequence, template proteins of similar folds are retrieved from the PDB library by LOMETS (LOcal MEta Threading Server)
- *Iterative Monte-Carlo simulation for structural assembly*: continuous fragments expunged from the PDB templates are assembled into full-length models using Monte Carlo simulations, and loops are constructed using *ab initio* structure modelling.
- *Selection of model and refinement*: simulation is performed to remove steric clashes between atoms of the amino acid residues. The global topology of the constructed model is improved, and the final structures are built by optimising the hydrogen bonds.
- *Function annotation*: function is inferred by comparing the structural models with known proteins.

The final output of I-TASSER includes the top ten structural models for bovine TLR along with the top ten alignments and top ten PDB structures used in model building and are the closest to the modelled structure.

The PatchDock [56] and FireDock [57] servers were used for performing molecular docking analysis of the TLRs and bovine TB vaccine. The PatchDock server computes the docking transformations of the receptors and ligands based on their molecular shape complementarity. The results of PatchDock were then presented to FireDock for further refinement of the docking results and predicting the global binding energy of receptor (TLR) and ligand (bovine vaccine). The interaction of the docked complex (TLRs and bovine TB vaccine) was interpreted by PDBsum [58].

### 2.3 Framework for Drug target identification of *Mycobacterium bovis* using subtractive proteomic analysis

This section details the approach used to identify drug targets (Figure 2).

#### 2.3.1 Exclusion of proteins homologous with the host proteome

The conserved proteins identified by subtractive reverse vaccinology analysis, present among 11 strains of *Mycobacterium bovis*, were first tested for homology with the host proteome. The drugs targeting homologous proteins may cause side effects or hypersensitive reactions in the host. To identify homologous proteins between *Mycobacterium bovis* and the host (cattle), BLASTp was used [59]. BLASTp was performed using a non-redundant protein sequence database with a threshold E-value of e10-4. *Bos* (taxid:9903), *Bos tauurus* (taxid:9913) and *Bos indicus* (taxid:9915) were used as reference organisms for performing sequence similarity. BLASTp performs local sequence alignment and measures sequence similarities based on maximum segment pairs (MSP) [24]. The results of BLASTp consist of homologous and non-homologous sequences. The proteins with sequence identity and a bit score of more than 30% and 100, respectively, were excluded from the study.

#### 2.3.2 Eliminating proteins involved in common metabolic pathways with the host

Comparative metabolic pathway analysis was carried out to identify proteins involved in metabolic pathways common in cattle and *Mycobacterium bovis.* The Kyoto Encyclopaedia of Gene and Genome (KEGG) [60] was used for performing this step. A manual comparison was made, and the proteins involved in the common pathways were removed.

#### 2.3.3 Unique metabolic pathway analysis

A drug target should be uniquely present in the pathogen to treat an infectious disease. The selection of proteins involved in the specific pathways of *Mycobacterium bovis* was made using KEGG automatic annotation server (KAAS) [61]. BLAST was selected for performing the search in KAAS. *Mycobacterium bovis* AF2122/97 was chosen as a reference from the organism list. A manual investigation was carried out to select proteins unique to the *Mycobacterium bovis* AF2122/97 strain, while the remaining proteins were excluded from the study.

#### 2.3.4 Retrieval of the essential proteins of *Mycobacterium bovis*

In this step, the identification of proteins that play a vital role in biological processes and functions, such as physiology, metabolism and developmental processes within *Mycobacterium bovis* was done. These proteins help in the regular functioning and survival of the disease-causing *Mycobacterium bovis*. For the identification of essential proteins, the Database of Essential Genes (DEG) was used. The database contains information about essential genes and their proteins obtained from different experimental methods [62]. For a query sequence, a similarity search was performed against the essential proteins present in DEG with the help of BLASTp. The parameters for the search were set as: e-value 10-50; sequence identity more than 30%; bit score >100 and query coverage 100%.

#### 2.3.5 Prediction of the role of proteins in virulence

The proteome of *Mycobacterium bovis* consists of proteins involved in virulence, the progression of bovine TB and escaping the host’s immune response [4]. Virulent proteins play a crucial role in infectious pathways, such as adherence, colonisation, invasion, and help escape the host’s immune response. Drugs targeting the pathogen’s virulent pathway would help eliminate the disease from the host body. Thus, there is a need to identify proteins that play a significant role in causing the infection in the host cell. Two tools were used for predicting virulent proteins: VirulentPred, an SVM-based method [37], and the MP3 tool, which uses SVM and HMM [38]. Proteins predicted virulent by both methods were selected for further analysis.

#### 2.3.6 Codon adaptation index

The codon adaptation index (CAI) is a measure of synonymous codon usage bias [63]. CAI predicts the translational efficiency and level of expression of a gene. The EMBOSS server was used to calculate the codon adaptation index for predicting the expression level of the selected gene targets of *Mycobacterium bovis* inside the host body. Proteins that are highly expressed are considered potential drug targets. The nucleotide sequences of the selected proteins were obtained from the NCBI Gene database to calculate CAI. The gene sequences were then submitted to EMBOSS server to predict each gene’s CAI. The genes with a CAI score of more than 0.7 were selected for further analysis.

#### 2.3.7 Analysis of physiochemical properties of selected drug targets

ProtParam was used for computing various physiochemical properties of the chosen drug targets in *Mycobacterium bovis*. The properties include molecular weight (MW), isoelectric point of a protein (pI), amino acid composition, extinction coefficient [48], instability index (II), estimated half-life [64], aliphatic index [49] and the grand average of hydropathicity (GRAVY) [47]. A potential drug target protein should have a molecular weight lower than 100 kDa. Isoelectric point is the pH of the solution at which the net electrical charge of the protein becomes zero [65]. The instability index determines the stable nature of the protein in a laboratory test tube. A protein with an instability index lower than 40 is a potential drug target. The extinction coefficient predicts the absorption of light at a particular wavelength by the target protein and helps determine the protein concentration. The GRAVY score determines the hydrophilic or hydrophobic nature of a protein.

#### 2.3.8 Structural homology analysis

The selected proteins were then analysed for the availability of their crystallographic X-ray structure in the PDB database. For this, BLASTp was performed using a PDB database with a threshold E-value of e10-6. The results of BLASTp consist of homologous and non-homologous sequences. The proteins with sequence identity and a bit score of more than 30% and 100, respectively, were selected in the study. The proteins with no template structure available for generating the 3D structural model were eliminated.

#### 2.3.9 Prediction of druggability potency of the selected proteins

Druggability is the ability of a protein to bind to a drug molecule. The druggability of the selected proteins was predicted by performing a sequence similarity search in the DrugBank database. DrugBank is a trustworthy database providing information on the available drugs and their drug targets [66]. DrugBank currently contains information on 9591 drugs with 2037 FDA-approved drugs (Food and Drug Administration). Over 6000 experimental drug entries are present in DrugBank. The e-value for the similarity search was set to 10-20. The drug target proteins for whom homologous proteins are present in the DrugBank database are druggable proteins. The protein with no hits found against any protein sequence is a novel drug target.

## 3. Results

### 3.1 Identification of conserved proteins across 11 strains of *Mycobacterium bovis*

First, complete proteome sequences of 11 strains of *Mycobacterium bovis* that have been completely sequenced were downloaded using the NCBI Genome FTP site. *Mycobacterium bovis* AF2122/97 with 3988 proteins present in its proteome was used as a reference strain. A standalone BLAST was then used for a proteome comparison of 11 strains, and the results were stored in Excel files. Manual comparison of the results revealed that 1163 proteins were conserved with more than 99% sequence similarity and 100% query coverage among the 11 strains of *Mycobacterium bovis* (Supplementary Table S1).

### 3.2 Developing an epitope-based bovine TB vaccine

The use of BCG in cattle has not provided sufficient protection. Several attempts have been made in the past to develop a live-attenuated or heat-killed vaccine against bovine TB in cattle. This study developed a conceptual framework for designing an epitope-based bovine TB vaccine containing B-cell and T-cell (HTL and CTL) epitopes that could elicit a humoral and cell-mediated immune response.

#### 3.2.1 Identification of nine antigenic proteins through the subtractive reverse vaccinology approach

To develop an effective bovine TB vaccine, a subtractive reverse vaccinology approach was implemented to identify surface-exposed antigenic proteins from the proteome of *Mycobacterium bovis* AF2122/97 with critical functional features, such as membrane-spanning regions, signal peptides, lipoprotein signatures, adhesion probability and non-homology to host. First, the subcellular localization of each conserved protein was predicted. The proteins in the cell membrane, cell wall and extracellular space are engaged in membrane integrity and permeability, efflux mechanisms and active transport of molecules; thus, they are considered suitable vaccine targets. Six bioinformatics tools (PSORTb v.3.0, CELLO, LocTree3, SOSUI, pLoc_bal-mGpos and GramLocEN) were utilized for SCL prediction. Of the 1163 conserved *Mycobacterium bovis* proteins, 273 proteins, commonly predicted by more than four SCL tools, were selected for further analysis. In the next step, proteins with more than one transmembrane α-helix were excluded from the study, as they are challenging to clone and purify in the laboratory. TMHMM server helped shortlist 142 proteins with 0 or 1 transmembrane α-helix. The presence of signal peptides and lipoprotein signatures within a protein makes it a vital antigenic and immunogenic vaccine target. The Sec (secretory) and TAT signal peptides are ubiquitous protein-sorting signals that help translocate a protein across the cell membrane in *Mycobacterium bovis*. For the prediction of secretory signal peptides, the SecretomeP and SignalP 4.1 servers were used. A total of 43 conserved and surface-exposed proteins showed the presence of Sec signal peptides. Next, PRED-TAT was used to identify tat signal peptides, and 12 proteins were involved in the TAT pathway. Thus, out of 142 proteins, 55 proteins (Sec and TAT proteins) were engaged in the secretory pathway of *Mycobacterium bovis*. Lipoproteins, essential sets of membrane proteins performing vital functions in *Mycobacterium bovis,* were predicted using PRED-LIPO. Twenty-eight lipoproteins were identified from 142 proteins. A total of 83 proteins (55 secretory proteins and 28 lipoproteins) were carefully chosen for further analysis.

A protein that can activate an immune response against pathogenic bacteria without causing considerable side effects inside the host body is considered a potential vaccine target. The VaxiJen, VirulentPred and MP3 servers were used to determine the antigenicity of 83 proteins. Concordance analysis of the three antigen servers identified 21 highly antigenic proteins that can initiate a robust immune response with or without side effects. Thus, the allergenic nature of the selected antigens was determined. Algpred was used to predict the allergenicity of the antigenic proteins. From 21 antigenic proteins, 18 proteins were discovered to be non-allergenic. These 18 conserved, surface-exposed, antigenic and non-allergenic proteins were further analysed for adhesion probability. Adhesin proteins help in the attachment of *Mycobacterium bovis* to the host’s cell-surface receptors. Hence, SPAAN was used to identify adhesin proteins among the shortlisted proteins. Of the 18 proteins, only nine had a strong adhesion probability score of 0.5 or above.

The last subtractive reverse vaccinology step involves studying the homology between the host (cattle) and pathogen (*Mycobacterium bovis*). Homologous proteins can initiate hypersensitive reactions inside the host body, causing severe health problems or side effects. For homology analysis, BLASTp was used for determining sequence similarity among host and pathogen. *Bos tauurus* (taxid:9913) and *Bos indicus* (taxid:9915) were selected as reference organisms for sequence similarity. No similarity was observed among the nine proteins of *Mycobacterium bovis* and cattle. Finally, a total of nine conserved, membrane-spanning, antigenic and non-allergic proteins were selected with the help of a reverse vaccinology approach from the proteome of *Mycobacterium bovis,* as shown in Table 1.

**Table 1:** List of nine conserved, surface-exposed, antigenic and non-allergic proteins identified by the reverse vaccinology approach. Column 1-accession number of protein, column 2-the function of protein, columns 3-10 show the results of the reverse vaccinology process for the shortlisted proteins

| Accession number | Function | Prediction of signal peptide | TMHMM | MP3 | VirulentPr ed | VaxiJe n | VaxiJe n Score | AlgPre d | Adhesion probability |
| --- | --- | --- | --- | --- | --- | --- | --- | --- | --- |
| YP_009357464.1 | SECRETED ANTIGEN 85-C FBPC (85C) (ANTIGEN 85 COMPLEX C) (AG58C) (MYCOLYL TRANSFERASE 85C) (FIBRONECTIN-BINDING PROTEIN C) | Tat signal peptide predicted | Pred Helix=1 | Pathogenic | Virulent | Probabl e Antigen | 0.4402 | Non-allergen | 0.812 |
| YP_009358272.1 | ppe family protein ppe14 | Sec signal peptide predicted | Pred Helix=0 | Pathogenic | Virulent | Probabl e Antigen | 0.6802 | Non-allergen | 0.592 |
| YP_009359011.1 | pe family protein pe17 | Sec signal peptide predicted | Pred Helix=0 | Pathogenic | Virulent | Probabl e Antigen | 0.4152 | Non-allergen | 0.78 |
| YP_009359070.1 | ppe family protein ppe23 | Sec signal peptide predicted | Pred Helix=0 | Pathogenic | Virulent | Probabl e Antigen | 0.4637 | Non-allergen | 0.752 |
| YP_009361194.1 | SECRETED MPT51/MPB51 ANTIGEN PROTEIN FBPD (MPT51/MPB51 ANTIGEN 85 COMPLEX C) (AG58C) (MYCOLYL TRANSFERASE 85C) (FIBRONECTIN-BINDING PROTEIN C) (85C) | Sec signal peptide predicted | Pred Helix=1 | Pathogenic | Virulent | Probabl e Antigen | 0.4515 | Non-allergen | 0.747 |
| YP_009361195.1 | SECRETED ANTIGEN 85-A FBPA (MYCOLYL TRANSFERASE 85A) (FIBRONECTIN-BINDING PROTEIN A) (ANTIGEN 85 COMPLEX A) | Tat signal peptide predicted | Pred Helix=1 | Pathogenic | Virulent | Probabl e Antigen | 0.5259 | Non-allergen | 0.77 |
| YP_009358321.1 | POSSIBLE LIPOPROTEIN LPRP | Lipoprotein | Pred Helix=0 | Pathogenic | Virulent | Probabl e Antigen | 0.5052 | Non-allergen | 0.598 |
| YP_009358349.1 | POSSIBLE CONSERVED EXPORTED PROTEIN | Sec signal peptide predicted | Pred Helix=1 | Pathogenic | Virulent | Probabl e Antigen | 0.5621 | Non-allergen | 0.757 |
| YP_009361095.1 | CONSERVED HYPOTHETICAL PROTEIN | Sec signal peptide predicted | Pred Helix=0 | Pathogenic | Virulent | Probable Antigen | 0.4189 | Non-allergen | 0.619 |

#### 3.2.2 Prediction of B-cell and T-cell epitopes from the nine conserved antigens using an immunoinformatics approach

An ideal prophylactic bovine TB vaccine should stimulate a robust immune response by generating memory cells to eliminate the infection in the future. For this, there is a need to identify immunodominant B-cell and T-cell epitopes that would initiate humoral and cellular immunity inside the host. Hence, immunoinformatic analysis was performed to screen the effective B-cell and T-cell epitopes required to design an epitope-based bovine TB vaccine.

##### 3.2.2.1 Prediction of B-cell epitopes

B-cell epitopes help generate a humoral or antibody-mediated immune response in the host’s body. ABCpred was utilized for identifying B-cell epitopes from the nine antigenic proteins of *Mycobacterium bovis*. The length of the B-cell epitope was set to 20 amino acid residues. A total of 251 B-cell epitopes were found from the nine antigenic proteins. The immunogenic nature of B-cell epitopes was identified using the VaxiJen server and ToxinPred. Supplementary Table S2 provides the results of immunoinformatics analysis performed on these nine bovine TB antigens. Of the 251 epitopes, 80 B-cell epitopes were found to be antigenic and non-toxic. Next, the identification of non-allergenic B-cell epitopes was performed with the help of the ALLERTOP 2.0 server. A total of 60 non-allergenic epitopes were shortlisted for further analysis of their hydrophilicity. The hydrophilic residues on the surface of antigenic proteins facilitate quick interactions with the host’s immune cells. The GRAVY score was calculated using the ProtParam server. In the ProtParam analysis, 16 B-cell epitopes were discovered to have negative GRAVY scores. No potential B-cell epitopes were found in the YP_009359011.1 and YP_009359070.1 proteins. The 16 B-cell epitopes were selected for constructing the bovine TB vaccine (Table 2).

**Table 2:**
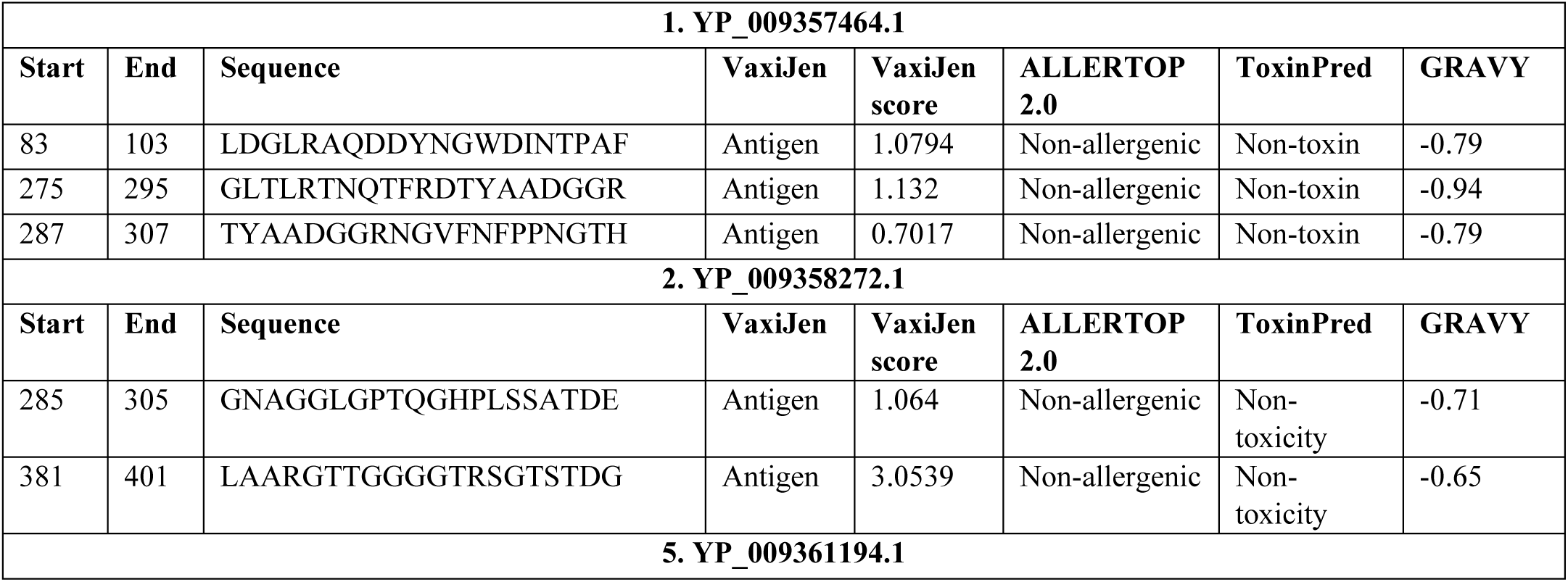
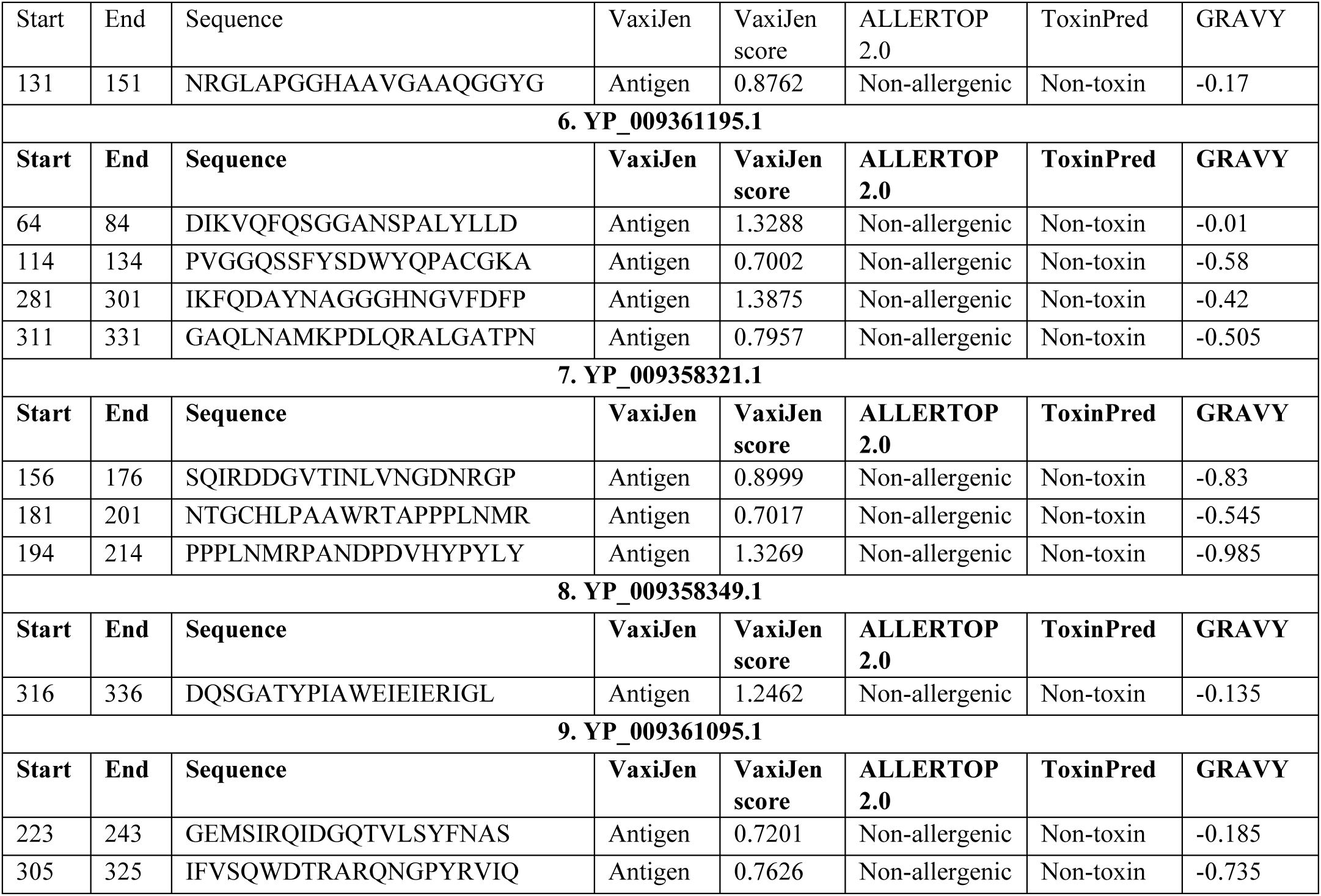
Shortlist of 16 B-cell epitopes from *Mycobacterium bovis* antigens required for constructing an epitope-based vaccine for bovine tuberculosis.

##### 3.2.2.2 Prediction of MHC class I restricted (CTL) T-cell epitopes

The antigens of *Mycobacterium bovis* are engulfed by the antigen-presenting cells and then fragmented into smaller antigenic peptides called epitopes. These epitopes are then presented to T-cell receptors (TCR) on the surface of T-cells through cell-surface attached MHC molecules. The epitopes binding to MHC class I molecules are termed cytotoxic T-cell (CTL) epitopes, or MHC class I restricted T-cell epitopes, and play an essential role in cellular immunity. For predicting MHC class I-restricted T-cell epitopes, the following bovine MHC class I HLA alleles were used: BoLA-D18.4, BoLA-AW10, BoLA-JSP.1, BoLA-HD6, BoLA-T2a, BoLA-T2b and BoLA-T2c. IEDB MHC-I and the NetMHCpan 4.1 server were used for predicting CTL epitopes. The length of CTL epitopes was set to nine amino acid residues. Both servers commonly predicted 78 CTL epitopes.

To validate the CTL epitopes’ ability to initiate a cellular immune response, antigenicity and toxicity analysis were completed using VaxiJen and Toxin Pred. Out of 78 CTL epitopes, only 24 epitopes were highly immunogenic and non-toxic. These 24 epitopes were then considered for allergenicity prediction using ALLERTOP 2.0. Three CTL epitopes found to trigger allergenic reactions inside the host body were eliminated. A hydrophilic score calculation was then undertaken using the ProtParam server. Epitopes with positive GRAVY scores were excluded from the study. Eight CTL epitopes, hydrophilic in nature with a negative GRAVY score, were chosen for constructing an epitope-based TB vaccine (Table 3). YP_009357464.1, YP_009359011.1, YP_009361195.1 and YP_009358349.1 antigens had no potential immunodominant CTL epitopes. Supplementary Table S3 shows the results of antigenicity, toxicity, allergenicity and hydrophilicity.

**Table 3:**
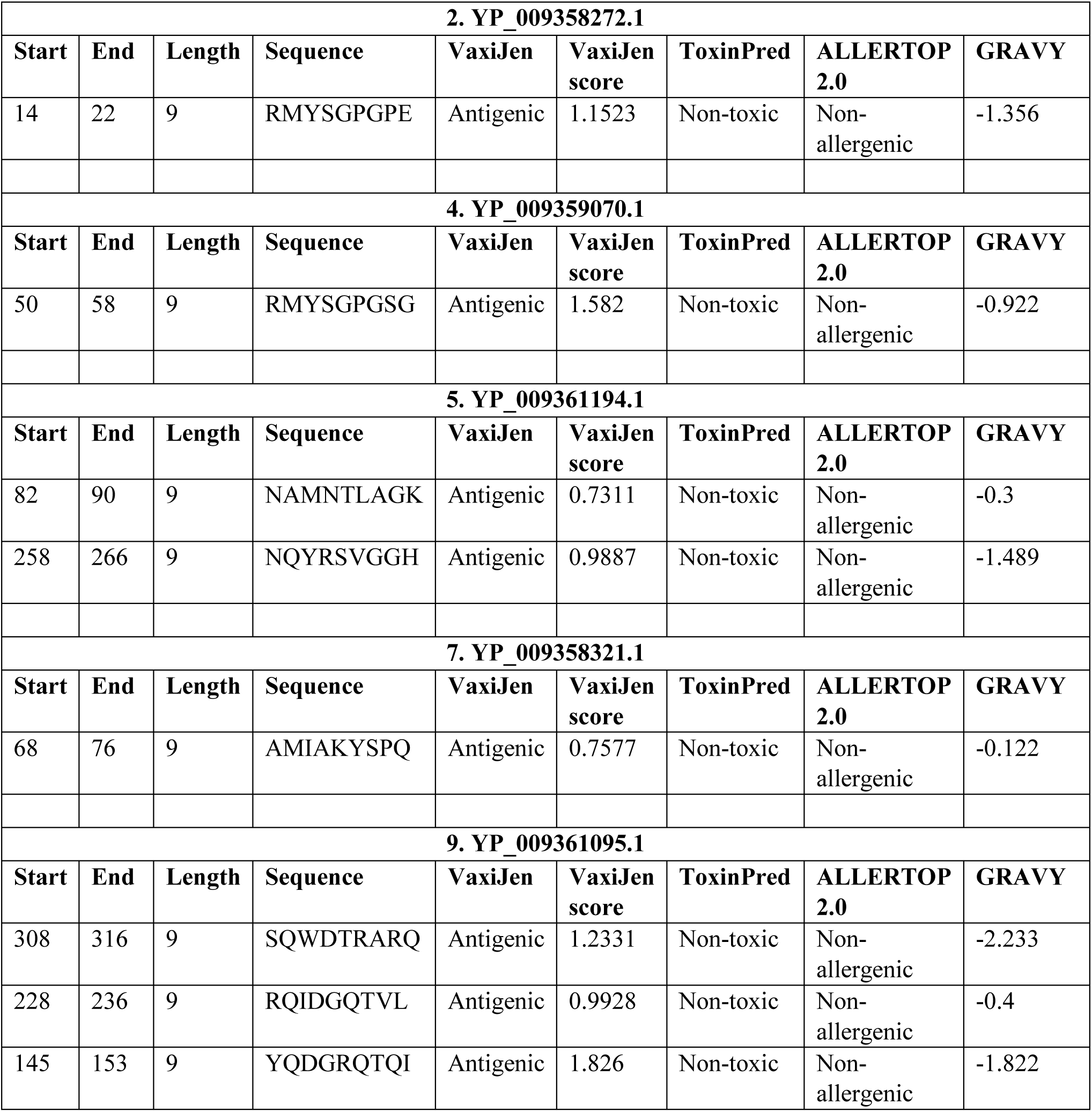
The eight shortlisted CTL epitopes from *Mycobacterium bovis* antigens required for constructing an epitope-based vaccine for bovine tuberculosis.

##### 3.2.2.3 Prediction of MHC class II-restricted (HTL) T-cell epitopes

The epitopes presented by MHC class II molecules are called helper T-cell epitopes (HTL epitopes) or MHC class II-restricted T-cell epitopes. The IEDB MHC II server was used to predict 15-mer HTL epitopes of *Mycobacterium bovis* using the consensus prediction method. HLA alleles of the mouse model were used in this step as no information was available for bovine MHC class II molecules. The lower percentile rank indicates a higher affinity of an epitope towards the MHC molecule. Thus, the epitope with percentile ranks lower than 1.0 were chosen. 145 HTL epitopes were predicted from the nine antigens of *Mycobacterium bovis* using the IEDB MHC II server. Next, 83 immunogenic and non-toxic HTL epitopes were discovered from 145 epitopes with the help of VaxiJen and ToxinPred. The allergenicity of 83 HTL epitopes was then predicted using the ALLERTOP 2.0 server. Fifty-two non-allergenic HTL epitopes were found. Finally, only two HTL epitopes were selected after calculating the hydrophilicity of the selected 52 non-allergenic HTL epitopes (Table 4). Supplementary Table S4 shows the results of immunoinformatics analysis on the nine antigens of *Mycobacterium bovis*.

**Table 4:**
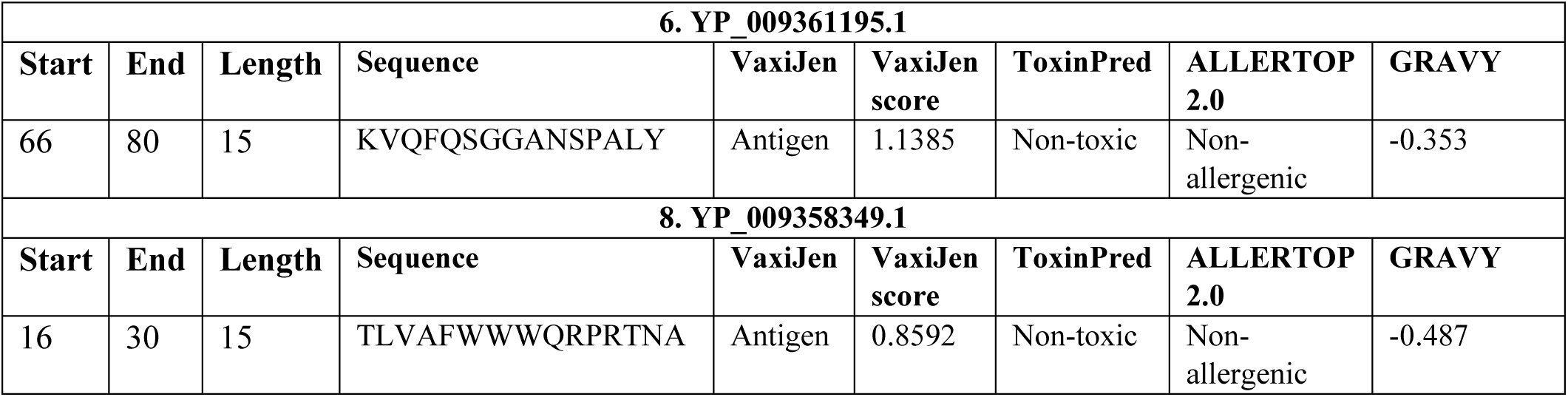
Two shortlisted HTL epitopes from *Mycobacterium bovis* antigens required for constructing an epitope-based vaccine for bovine tuberculosis.

#### 3.2.3 Designing the vaccine sequence and assessment of its properties

After selecting highly immunogenic and excluding cross-reactive epitopes, 26 epitopes (HTL epitopes-2, CTL epitopes-8 and B-cell epitopes-16) were shortlisted from the nine antigenic proteins of *Mycobacterium bovis* (Supplementary Table S5). The selected epitopes would evoke a potent cellular and humoral immune response inside the host body. The sequence of the bovine TB vaccine was designed using a simple strategy. First, the adjuvant 50s ribosomal protein L7/L12 at the N-terminal was added to the vaccine protein sequence with the help of the EAAAK linker (Figure 3(i)). HTL, CTL and B-cell epitopes were joined using flexible linker AAY. With one adjuvant, one EAAAK linker, 25 AAY linkers and 26 epitopes (HTL epitopes-2, CTL epitopes-8 and B-cell epitopes-16), the final length of the bovine TB vaccine sequence was 632 amino acid residues (Figure 3(ii)).

**Figure 3:**
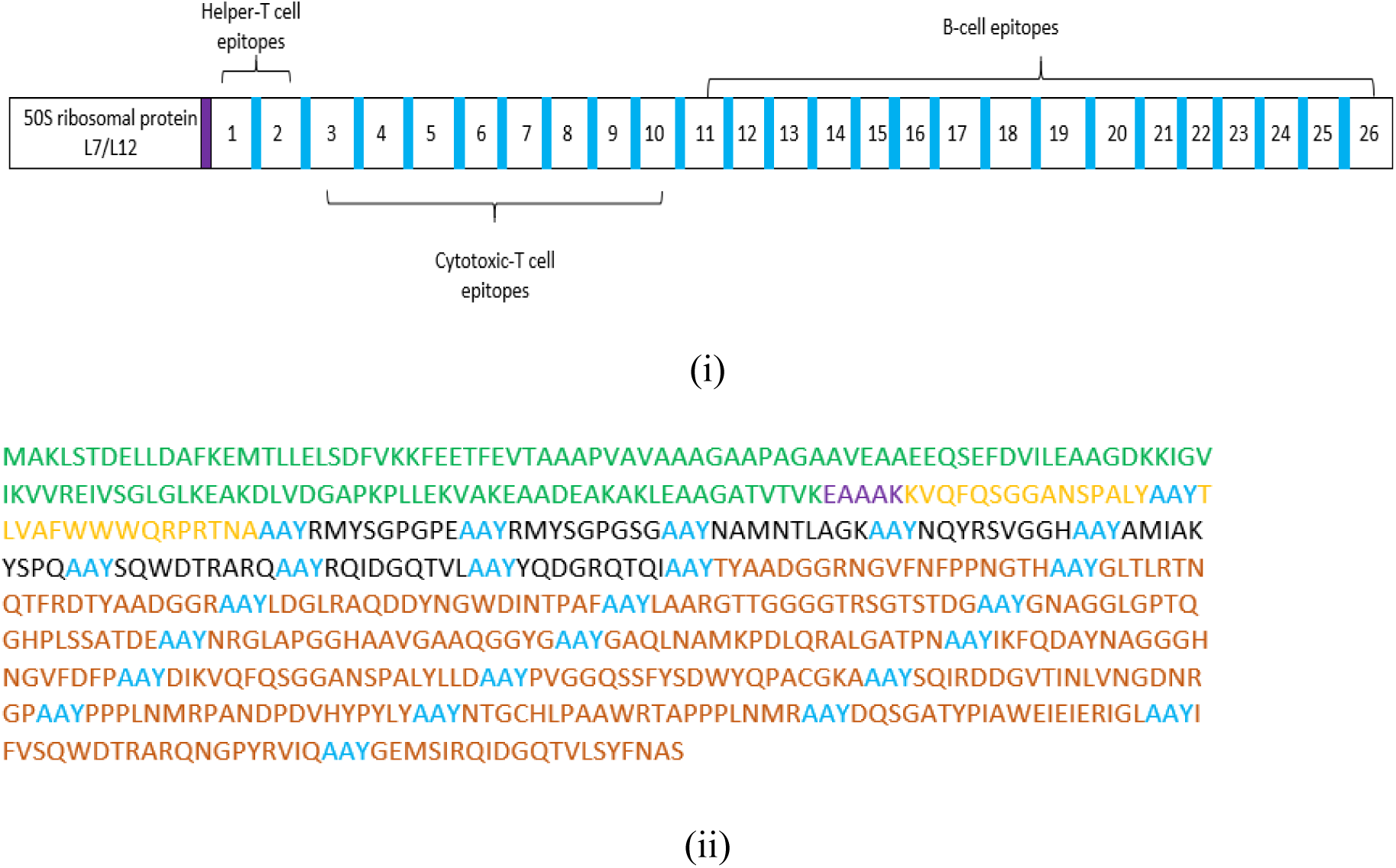
Epitope-based vaccine sequence construction scheme. (i) Schematic representation of bovine TB vaccine construct consisting of B-cell and T-cell epitopes joined together by flexible linkers and an adjuvant at the N-terminal of the vaccine protein, (ii) TB vaccine protein sequence. The adjuvant sequence is highlighted in green with flexible linkers in purple (EAAAK) and blue (AAY). HTL epitopes are highlighted in orange, with CTL epitopes in black and the B-cell epitopes in brown

Antigenicity, allergenicity and physiochemical properties were evaluated for the designed bovine TB vaccine sequence. The VaxiJen server was used to predict the antigenicity of the vaccine sequence. A high antigenic score of 0.82 showed the remarkable immunogenic potential of the epitope-based bovine TB vaccine. The AlgPred server suggested that the designed bovine TB vaccine was non-allergenic to the host (cattle).

Next, ProtParam was used to predict the physiochemical properties of the vaccine sequence. The vaccine sequence comprised 632 amino acid residues with a molecular weight of 66.85 kDa. Proteins with a molecular weight of less than 100 kDa are suitable vaccine candidates. The bovine TB vaccine construct contained 9071 atoms, and its molecular formula was C_2981_H_4509_N_823_O_912_S_12._ The theoretical pI was assessed to be 5.77. The estimated half-life was 30 hours in mammalian reticulocytes (*in-vitro*). The instability index with a value of 32.35 represented the stable nature of the bovine TB vaccine. The aliphatic index predicts the relative volume occupied by aliphatic side chains in a protein. A value for the aliphatic index greater than 30 indicates that the protein is thermodynamically stable. The higher the value of the aliphatic index (68.56 in this study), the higher the thermos-stability of the protein. The GRAVY score was found to be −0.302. The negative value indicates the hydrophilic nature of the vaccine. After evaluating all the properties, the designed bovine TB vaccine had all the attributes of a promising prophylactic vaccine candidate required for initiating a robust immune response inside the host.

#### 3.2.4 Constructing the structural model of the epitope-based bovine TB vaccine

The secondary structure of the bovine TB vaccine was evaluated using PSIPRED. According to PSIPRED, 239 amino acid residues were engaged in forming alpha-helices, constituting 37.82% of the overall bovine TB vaccine sequence. Ninety-nine amino acid residues participated in the formation of beta-strand (15.66%), and 294 residues formed the coils (46.52%) (Figure 4).

**Figure 4:**
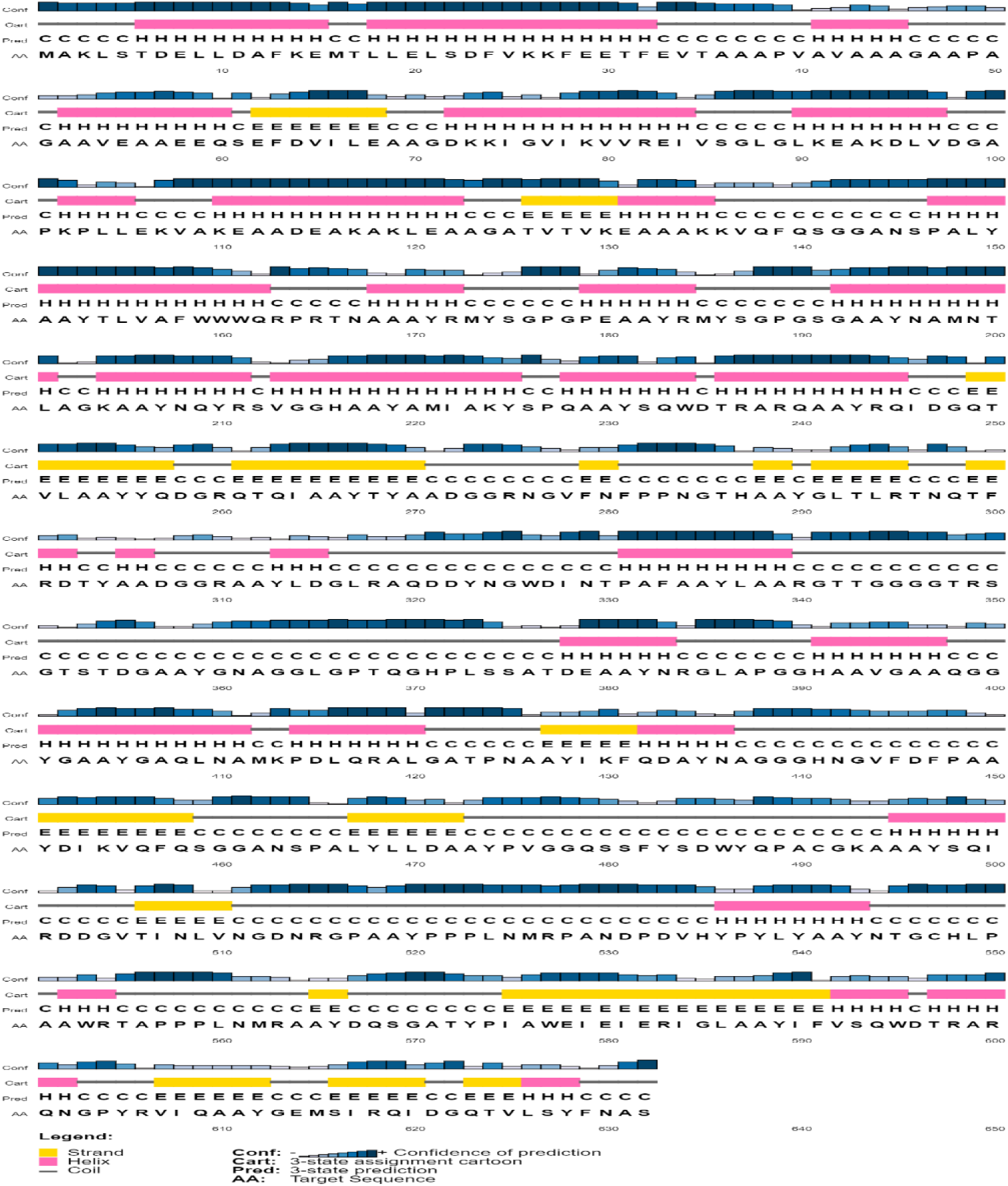
Secondary structure prediction for the bovine TB vaccine sequence using PSIPRED. The sequence comprised alpha-helices (37.82%), beta-strands (15.66%) and coils (46.52%)

The RaptorX server was used for constructing a tertiary structure model of the epitope-based bovine TB vaccine. It performed homology modelling using 6VHD_A PDB entry as a template for generating a structural model. All 632 amino acid residues of the vaccine protein were modelled with 8% in the disordered region. The constructed model was 39% exposed, 29% medium and 31% buried in folded conformation. The Ramachandran plot showed that 85.4% of amino acid residues were in the most favoured region. Thus, the GalaxyRefine server was used to refine the structural model to improve its quality. Figure 5(i) shows the 3D structural model of the bovine TB vaccine after refining with 91.1% amino acid residues in the most favoured region (Figure 5(ii)). Other parameters assessed after refinement were a GDT-HA score of 0.9568, MolProbidity 3.365 and RMSD 0.412.

**Figure 5:**
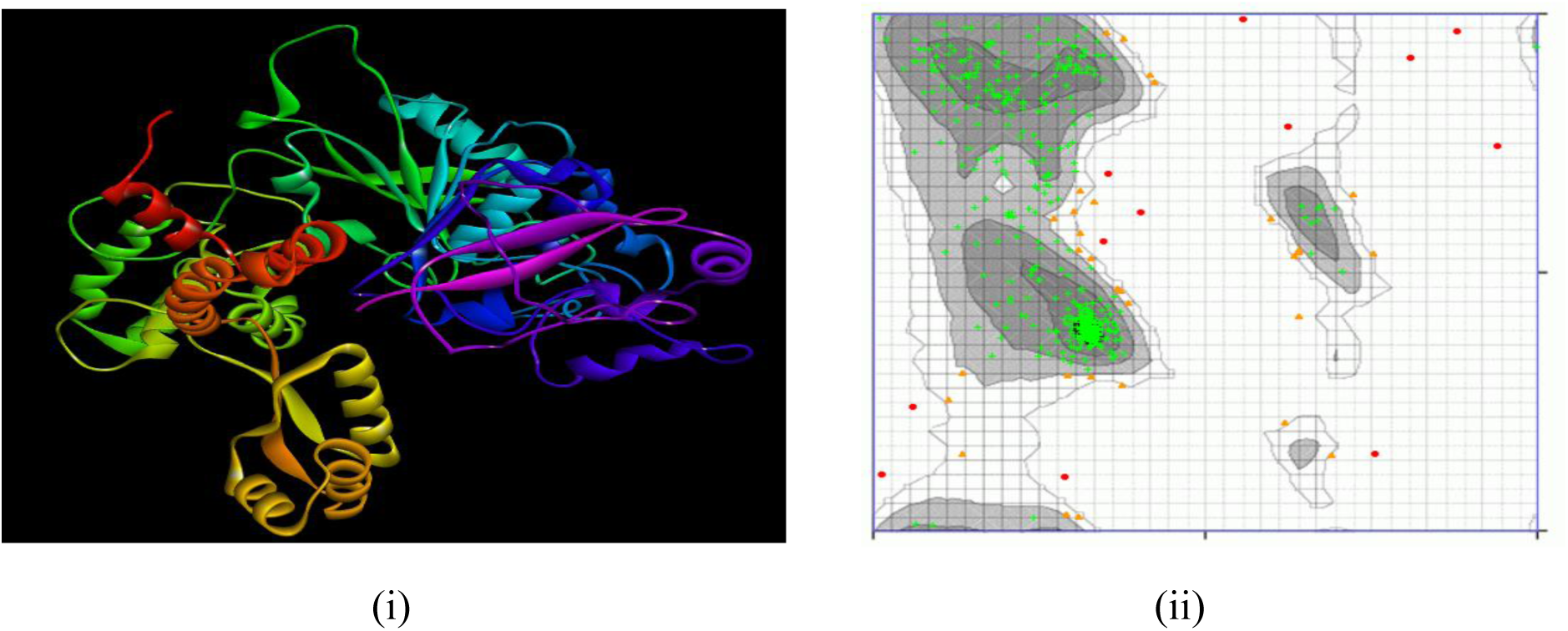
Refined structural model of bovine TB vaccine construct. (i) The 3D structure is coloured in rainbow colours, from violet to red (from N-terminus to C-terminus), and (ii) the Ramachandran plot of the refined structure shows 91.1% amino acid residues in the most favoured region (green crosses)

#### 3.2.5 Analysis of the interaction of bovine TB vaccine and toll-like receptors

TLRs interact with the pathogen/vaccine antigen to initiate the innate immune response. The stable interaction of a vaccine with receptors of host immune cells activates the immune system. To determine the ability of the designed bovine TB vaccine to generate an immune response, molecular docking analysis was performed. First, the structures of the bovine TLRs were constructed using the I-TASSER server. The protein sequences used for structure construction were Q95LA9, Q9GL65 and B5T278 for TLR-2, TLR-4 and TLR-6.

After the construction of the TLRs, PatchDock and FireDock were used to analyse the interactions between the bovine TB vaccine construct and the TLRs of cattle. The output of the PatchDock docking process displayed several docked transformations, which were further presented to FireDock for calculating the global binding energy. PatchDock found 1455 docked transformations for TLR-2 and the bovine TB vaccine docked complex (Table 5). The lowest binding energy of −55.15 kcal/mol was predicted for the 433^rd^ transformation of the docked complex. The global binding energies for the bovine TB vaccine docked with TLR-4 and TLR-6 were −61.78 kcal/mol and −53.89 kcal/mol, respectively. The docking results revealed that the bovine TB vaccine designed was adequately engaged in interaction with the selected TLRs (Figure 6). The docking interaction of TLRs and bovine TB vaccine was analysed using PDBsum. The interaction of TLR-2 and bovine TB vaccine showed 5 hydrogen bonds and 179 non-bonded contacts; 4 hydrogen bonds and 209 non-bonded contacts were examined TLR-4 and TB vaccine, and 7 hydrogen bonds, 1 salt bridge, and 169 non-bonded contacts were analysed for TLR-6 and bovine TB vaccine. The docking results revealed that the designed TB vaccine had substantial interaction with the selected TLRs and would accomplish the goal of initiating the immune response.

**Figure 6:**
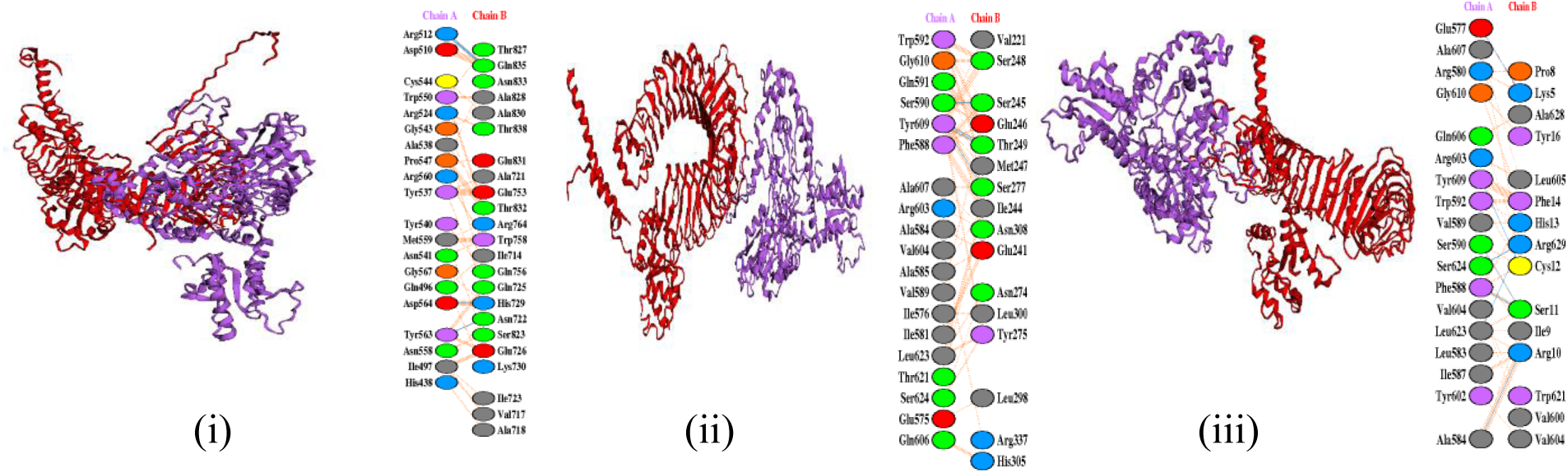
The docked complex of TLRs (red colour) with the TB vaccine construct (purple colour). TB vaccine construct (Chain A) docked with TLR (Chain B): (i) TLR-2, (ii) TLR-4 and (iii) TLR-6

**Table 5:** Molecular docking of bovine the TB vaccine and the toll-like receptors of the host (cattle). (aVdW-attractive Van der Waals energy, rVdW-repulsive Van der Waals energy and HB-hydrogen bonding)

| Docked Complex | Total number of docked transformations | Transformation with lowest binding energy | Global binding energy (kcal/mol) | aVdW (kcal/mol) | rVdW (kcal/mol) | HB (kcal/mol) |
| --- | --- | --- | --- | --- | --- | --- |
| Bovine TB vaccine-TLR2 | 1455 | 433rd | -55.15 | -39.38 | 26.12 | -4.1 |
| Bovine TB vaccine-TLR4 | 1398 | 37th | -61.78 | -39.51 | 22.19 | -4.7 |
| Bovine TB vaccine-TLR6 | 2911 | 12th | -53.89 | -36.96 | 20.78 | -5.21 |

### 3.3 Identification of nine potential drug candidates from the 1163 conserved proteins within 11 strains of *Mycobacterium bovis*

This part of the study aimed to identify potential drug candidates for bovine TB. Some crucial steps were performed to discover unique drug targets that are pivotal for the survival of *Mycobacterium bovis* and do not initiate hypersensitive reactions or side effects in the host. In this study, *Mycobacterium bovis* AF2122/97 was taken as a reference organism for research, and it has a total of 3988 protein sequences in its proteome. 1163 conserved proteins were identified within 11 strains of bovine TB bacteria through comparative proteomic analysis. Using the 1163 conserved proteins, subtractive proteomics was performed approach to identify drug targets that could further help investigate therapeutic drugs for treating bovine TB. Table 6 shows the summary of the results of the subtractive proteomic analysis

**Table 6:** Exploration of potential drug targets for bovine TB using the subtractive proteomic approach for *Mycobacterium bovis*.

| Steps | Result |
| --- | --- |
| Total proteins | 3988 |
| Conserved proteins | 1163 |
| Non-homologous proteins to cattle | 815 |
| Non-homologous host metabolic pathways | 735 |
| Proteins in unique metabolic pathways | 386 |
| Essential metabolic proteins | 195 |
| Virulent proteins | 29 |
| Highly expressed target proteins | 13 |
| Potential bovine TB drug target | 9 |

The homologous proteins present in *Mycobacterium bovis* and the host are not considered to be effective drug targets. Thus, to exclude homologous proteins, sequence similarity analysis was done using BLASTp. The protein sequence identities of 30% or more, i.e., bit scores above 100, were eliminated from the study. Out of 1163 conserved proteins, 348 homologous protein sequences were found. These 348 proteins were excluded, and the 815 non-homologous proteins were considered for further examination. An analysis to identify proteins involved in metabolic pathways common in cattle and Mycobacterium bovis was implemented to enhance the drug targets’ safety profile. Seven hundred and thirty-five non-homologous proteins were subsequently obtained that were not involved in metabolic pathways similar to the host’s metabolic system.

To determine the involvement of these 735 proteins in unique metabolic pathways, KEGG Automated Annotation Server (KAAS) was used. In the analysis, more than 50% of the non-homologous proteins were involved in distinctive pathways in *Mycobacterium bovis.* A total of 386 proteins were involved in the unique metabolic pathways of *Mycobacterium bovis*. The proteins retrieved were first classified as enzymatic or non-enzymatic with the help of enzyme classification (EC) numbers. Around 56.74% of the proteins were enzymatic, and 43.26% were non-enzymatic. In the KEGG database, the pathways were broadly classified into seven categories: metabolism, genetic information processing, environmental information processing, cellular processes, organism system, diseases and drugs. Out of 386 proteins, 80.05% (309 proteins) were involved in different metabolic pathways (Figure 7). For example, 60 proteins were involved in amino acid metabolism, followed by 48 in carbohydrate metabolism, 34 in energy metabolism, 18 in lipid metabolism, 11 in nucleotide metabolism and the remaining proteins were involved in the biosynthesis of secondary metabolites, cofactors and vitamins, terpenoids and polyketides. Approximately 26 proteins (14%) were involved in genetic and environmental information processing, 14 in cellular processes (3%), four in the organism’s system, three in diseases and four in drug categories (Figure 7). Supplementary Table S6 shows the classification of the 386 proteins based on the protein families involved in metabolism, genetic information processing, signalling mechanism, cellular processes, organism system, diseases and drugs.

**Figure 7:**
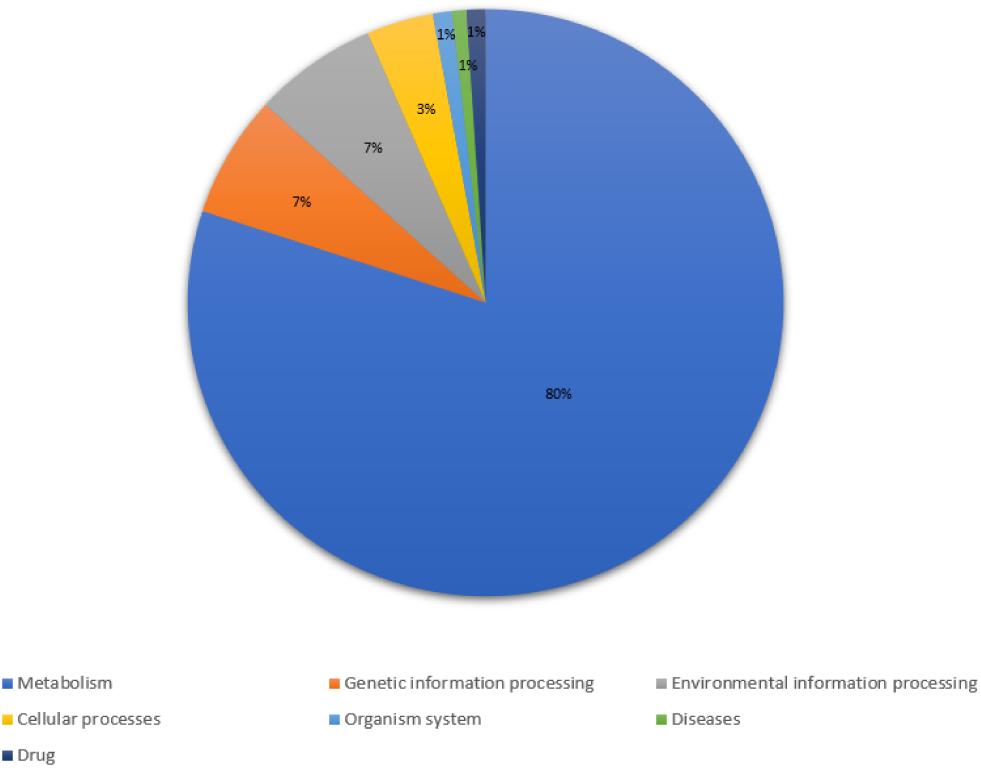
Percentage distribution of non-homologous proteins in seven pathways: metabolism, genetic information processing, environmental information processing, cellular processes, organism system, diseases and drugs

Identifying non-homologous proteins involved in unique metabolic pathways is not the sole criterion for selecting potential drug targets for bovine TB. A protein might be involved in more than two or three metabolic pathways, but it could be a non-essential protein for *Mycobacterium bovis* survival. Thus, it is crucial to target essential proteins that disrupt the normal functioning of bovine TB bacteria. Based on the criteria mentioned above, the DEG database was utilized to select proteins that are important for the survival of *Mycobacterium bovis*. Proteins showing good sequence similarity, 100% query coverage and a bit score of more than 100 with the laboratory-validated essential proteins present in DEG were considered essential. A total of 195 essential proteins were retrieved in this step.

Other crucial criteria for the identification of drug targets are virulence and pathogenicity. A non-homologous and essential protein involved in initiating the virulence mechanism in the host is considered a pivotal drug candidate. Targeting virulent proteins that help survive pathogenic bacteria inside the host would swiftly eliminate bovine TB disease. VirulentPred and MP3 tools were used to predict the virulence of essential proteins. Twenty-nine proteins, commonly predicted as virulent by both methods, were selected for further analysis. In the next step, the codon adaptation index (CAI) was calculated. CAI indicates the translational efficiency and level of expression of a gene. CAI helps choose highly expressed proteins from the set of selected essential and virulent proteins. The value of the CAI score ranges from 0 to 1 in the bacterial genome, and proteins with a CAI score of more than 0.7 are likely to be highly expressed in the *Mycobacterium bovis* genome. Of the 29 virulent proteins, only 13 proteins were found to have a CAI score greater than 0.7. Supplementary Table S7 provides information on the 13 proteins selected. The chosen proteins were further analysed for their physiochemical properties, structural homology and druggability properties.

ProtParam was used to predict the physiochemical properties of the selected 13 proteins of *Mycobacterium bovis* and these are shown in Table 7. The range of amino acid length was from 224-679 residues and the molecular weights varied from 21-72 kDa. All 13 proteins had a molecular weight lower than 100 kDa or 100000 Da, making them possible drug targets. Eight of the 13 proteins were stable, with an instability index lower than 40. The presence of dipeptides was the reason for the unstable nature of the remaining five proteins. The aliphatic index predicts the relative volume occupied by the aliphatic side chains in a protein. A value of aliphatic index greater than 30 indicates that the protein is thermodynamically stable. All 13 proteins had an aliphatic index greater than 30, marking the highly thermodynamically stable nature of the selected protein molecules. The GRAVY score predicts the hydropathy value, indicating a protein molecule’s hydrophilic or hydrophobic nature. Table 7 shows that six proteins had negative GRAVY scores indicating their hydrophilic nature. The remaining proteins had a GRAVY score below one, suggesting they are less hydrophobic molecules. Analysis of physiochemical properties showed that the selected 13 proteins have all the characteristics of potential drug targets.

**Table 7:** Physiochemical properties of the 13 non-homologous, essential, virulent and highly expressed proteins of *Mycobacterium bovis*.

| Protein Accession | Amino acids | MW (Dalton or Da) | pI | Extinction | Instability index | Aliphatic index | GRAVY |
| --- | --- | --- | --- | --- | --- | --- | --- |
| YP_009357346.1 | 491 | 51575.6 | 8.59 | 36120 | 32.66 | 87.17 | -0.085 |
| YP_009357462.1 | 455 | 49889.11 | 5.06 | 69455 | 38.05 | 88.62 | -0.185 |
| YP_009357464.1 | 340 | 36771.27 | 5.92 | 85830 | 49.34 | 68.97 | -0.253 |
| YP_009357822.1 | 369 | 38521 | 6.4 | 25690 | 27.64 | 99.95 | 0.133 |
| YP_009358286.1 | 304 | 32988.36 | 10.35 | 54430 | 41.3 | 129.51 | 0.868 |
| YP_009358342.1 | 504 | 54431.72 | 9.67 | 43430 | 50.18 | 102.38 | 0.006 |
| YP_009358652.1 | 325 | 35083.25 | 10.68 | 46410 | 27.19 | 120.34 | 0.517 |
| YP_009358831.1 | 591 | 62694.38 | 10.95 | 112535 | 36.24 | 121.4 | 0.631 |
| YP_009358851.1 | 241 | 24623.99 | 9.71 | 26930 | 31.48 | 81.87 | 0.033 |
| YP_009359528.1 | 679 | 72537 | 9.64 | 49975 | 43.17 | 79.47 | -0.346 |
| YP_009359962.1 | 224 | 21497.63 | 6.14 | 10095 | 26.06 | 98.35 | -0.021 |
| YP_009360863.1 | 449 | 49578.72 | 9.37 | 81500 | 38.34 | 102.92 | 0.461 |
| YP_009361152.1 | 234 | 25843.77 | 5.65 | 19940 | 52.93 | 104.49 | -0.059 |

A template in the PDB database makes a protein a suitable target for drug discovery and development. For this, structural similarity analysis was performed using BLASTp. The selected proteins of *Mycobacterium bovis* were investigated for structural homology. The proteins with a sequence identity and bit score of more than 30% and 100, respectively, were selected in the study (Table 8). Out of the 13 proteins, four proteins with no suitable template structure were eliminated from the study.

**Table 8:** Results of template search analysis of the selected nine potential drug targets.

| Protein Accession | Query cover | Sequence similarity | PDB ID |
| --- | --- | --- | --- |
| YP_009357346.1 | 93 | 100 | 3LO7_A |
| YP_009357462.1 | 100 | 99.78 | 4O7O_A |
| YP_009357464.1 | 86 | 100 | 4MQM_A |
| YP_009357822.1 | 100 | 100 | 5JZX_A |
| YP_009358342.1 | 30 | 72.26 | 6BLK_A |
| YP_009358851.1 | 87 | 100 | 3PBI_A |
| YP_009359528.1 | 82 | 99.82 | 6KGH_A |
| YP_009359962.1 | 95 | 32.57 | 2XF4_A |
| YP_009361152.1 | 95 | 54.67 | 5ED4_A |

Finally, the druggability potential of the nine shortlisted drug targets was predicted using DrugBank. Six target proteins were found to have significant hits in DrugBank with bit scores of more than 100 (Table 9). The six proteins (YP_009357346.1, YP_009357464.1, YP_009357822.1, YP_009358342.1, YP_009359528.1 and YP_009361152.1) with considerable similarity found in DrugBank, were termed druggable target proteins. The remaining three (YP_009357462.1, YP_009358851.1 and YP_009359962.1) proteins with no significant hits found were considered novel target proteins.

**Table 9:** Identified drugs, FDA approval or experimental compounds for them and their action on the six druggable proteins from the DrugBank.

| Protein Accession | Potential drugs | Drug group | Action |
| --- | --- | --- | --- |
| YP_009357346.1 | Cefepime, Cefiderocol | Approved | Inhibitor |
| YP_009357464.1 | Diethyl phosphonate | Experimental | Unknown |
| YP_009357822.1 | Flavin adenine dinucleotide | Approved | Unknown |
| YP_009358342.1 | Phosphoaminophosphonic acid-adenylate ester | Experimental | Unknown |
| YP_009359528.1 | Cefepime, Cefiderocol, Doripenem, Cefpodoxime | Approved | Inhibitor |
| YP_009361152.1 | Adenosine-5'-RP-alpha-thio-triphosphate | Experimental | Unknown |

The two FDA-approved drugs, Cefepime and Cefiderocol, target the same proteins, having accession numbers YP_009357346.1 and YP_009359528.1. Cefepime and Cefiderocol have the same mode of action as beta-lactam antibiotics, such as penicillin. They disrupt the synthesis of peptidoglycan in the bacterial cell wall, thus, weakening the structural integrity of the pathogenic bacteria. Cefpodoxime was found to be vet-approved with the potential to target the YP_009359528.1 protein by inhibiting the synthesis of peptidoglycan, deteriorating the bacterial cell wall. Cefpodoxime, flavin adenine dinucleotide approved compound, could target YP_009357822.1 protein, but the mechanism of compound action is not yet established. Experimental compounds were found for YP_009357464.1, YP_009358342.1 and YP_009361152.1, but no mechanism of action is known.

In the present study, nine proteins were selected as potential broad-spectrum drug targets (YP_009357346.1, YP_009357462.1, YP_009357464.1, YP_009357822.1, YP_009358342.1, YP_009358851.1, YP_009359528.1, YP_009359962.1 and YP_009361152.1) for bovine TB treatment that minimises the chances of drug resistance. YP_009357346.1 was a probable penicillin-binding protein having 491 amino acid residues in its sequence. This protein was involved in peptidoglycan biosynthesis, thus, maintaining the cell wall and cellular structure of *Mycobacterium bovis*. The disruption of this protein by a drug would cause structural irregularity and loss of cell wall integrity, leading to the weakening of bacterial cells. Two potential drugs, Cefepime and Cefiderocol, were found to have the capability to disrupt the YP_009357346.1 protein. YP_009357464.1 is present in the cell wall of *Mycobacterium bovis*. This is involved in lipid metabolism for the biosynthesis of arabinogalactan. Arabinogalactan is an important cell wall constituent and strengthens the cell wall of *Mycobacterium bovis*. Inhibition of the biosynthesis of arabinogalactan can make the bacteria fragile. DrugBank identified diethyl phosphonate as a potential experimental compound for targeting this protein.

YP_009357822.1, present in the cytoplasm, is involved in amino sugar and nucleotide sugar metabolism with a significant role in bacterial cell division. Based on its function, targeting this protein would interfere in the cell division of *Mycobacterium bovis.* DrugBank identified the flavin adenine dinucleotide compound with the potential to target YP_009357822.1 protein, but the compound’s action is unknown. YP_009358342.1, found in the cell membrane of *Mycobacterium bovis,* is involved in the environment information processing pathway. This two-component system in mycobacteria plays a crucial role in the signal transduction mechanism required to continue the bacterial infection. Attacking this protein with drugs would lead to halting the signalling mechanism, thus, containing the spread of bacterial infection. The phosphoaminophosphonic acid-adenylate ester experimental compound could target YP_009358342.1 protein, but the mechanism of the compound’s action is not yet established.

YP_009359528.1, a probable penicillin-binding membrane protein, pbpB, present in the cell wall, is involved in peptidoglycan biosynthesis. Four potential FDA-approved drugs were identified for this drug target: Cefepime, Cefiderocol, Doripenem and Cefpodoxime. The role of all four drugs is inhibition of the synthesis of peptidoglycans. Cefpodoxime was found to be a vet-approved drug. YP_009361152.1, a two-component system protein, was found in the cytoplasm of *Mycobacterium bovis*. This protein is involved in the environment information processing pathway. Adenosine-5’-RP-alpha-thio-triphosphate was found to have the potential for attacking the YP_009361152.1 protein.

The three proteins (YP_009357462.1, YP_009358851.1 and YP_009359962.1) with no significant hits found were considered to be novel target proteins. The YP_009357462.1 protein was found to be involved in carbohydrate metabolism and present in the cytoplasm of *Mycobacterium bovis*. The primary function of this protein is to transfer phosphorus-containing groups in starch and sucrose metabolism. Targeting carbohydrate metabolism usually induces a stress response in the cell leading to the death of bacteria. YP_009358851.1 is involved in peptidoglycan biosynthesis and degradation. An effective drug with an inhibitory mechanism would disrupt the synthesis of peptidoglycan. YP_009359962.1 present in the cytoplasm is involved in carbohydrate metabolism, specifically, pyruvate metabolism. This protein benefits the bacterial cell during stress conditions. Thus, YP_009359962.1 is a potential target for enhancing stress inside the bacterial cell. The nine proteins selected are highly conserved, unique to *Mycobacterium bovis*, and involved in essential and virulent pathways. These nine proteins can therefore be confidently used for drug discovery and development against *Mycobacterium bovis*.

## 4. Discussion and Conclusion

Bovine TB is an infectious disease caused by the pathogenic bacteria, *Mycobacterium bovis.* It has become of global concern over the last two decades. Cattle are considered the main reservoir of bovine TB compared to other domestic livestock. Bovine TB is a zoonotic disease that can infect humans by inhaling aerosols or ingesting unpasteurised milk. Thus, the zoonotic potential of bovine TB is raising health concerns for the public. Out of 10 million human incidences in 2019, WHO estimated 0.14 million cases were zoonotic TB caused by *Mycobacterium bovis*, with 11,400 human deaths [12]. Bovine TB is more common in less developed and developing countries. The disease can lead to a severe economic crisis because of significant livestock losses and subsequent trade restrictions. Isolating and slaughtering the infected animals are undertaken to reduce transmission among other animals in the herd. The isolation of infected animals is not an option in hugely populated or low-income countries. Currently, there is no effective treatment available for bovine tuberculosis due to its infectious nature and the drug resistance of *Mycobacterium bovis*. The available treatment for bovine tuberculosis mainly depends on the health status of the infected animal species. Antibiotic therapy can only be given to the animal species living in captivity, as it is complicated for herd or free-grazing animals. The BCG vaccine is another option for treating disease but shows limited efficacy in cattle. There is no permanent treatment and prevention for bovine tuberculosis.

The present study provides a potential solution for bovine TB treatment by identifying therapeutic targets. With this aim, an effective epitope-based vaccine for bovine TB was proposed. The availability of the complete proteome of *Mycobacterium bovis* facilitates the use of many computational approaches using bioinformatics tools in such endeavours. The present study incorporates different branches of bioinformatics, such as subtractive reverse vaccinology, immunoinformatics and structural vaccinology, to identify potential vaccine candidates and design an epitope-based bovine TB vaccine. The strategy employed in this study is an efficient way of enriching the potential bovine TB vaccine targets and selecting the highly immunodominant epitopes from the conserved and surface-exposed antigenic target proteins of *Mycobacterium bovis*. 11 strains of *Mycobacterium bovis* were shortlisted to cover their diversity and to determine the conserved proteins among them. *Mycobacterium bovis* AF2122/97 was used as a reference proteome for comparative analysis. Using standalone BLAST, 1163 proteins were discovered to be conserved among 11 strains of *Mycobacterium bovis*. After identifying conserved proteins, a reductionist reverse vaccinology process was performed to identify non-homologous, surface-exposed, antigenic and non-allergenic proteins having unique characteristics, such as signal peptides, membrane-spanning regions, lipoprotein signatures and adhesion probabilities. Out of 1163 conserved proteins, nine conserved, membrane-spanning, antigenic and non-allergic proteins were selected from the reverse vaccinology approach. Next, extensive immunoinformatic analysis was performed, and 26 epitopes (HTL epitopes-2, CTL epitopes-8 and B-cell epitopes-16) were shortlisted from the nine antigens of *Mycobacterium bovis*. These 26 epitopes were highly immunogenic, non-toxic and non-allergenic to the host.

The epitopes selected were used for designing the vaccine sequence. The sequence of the bovine TB vaccine was created using a simple strategy. A 50s ribosomal protein L7/L12 adjuvant was attached at the N-terminal with the help of the EAAAY linker followed by HTL, CTL and B-cell epitopes bound together, using the flexible linker, AAY. Assessment of antigenicity, allergenicity and the physiochemical properties of the bovine TB vaccine showed that the vaccine sequence designed had all the qualities of a potential vaccine that would generate an effective immune response inside the host. A model of the bovine TB vaccine was constructed using RaptorX. Structural analysis of the epitope-based bovine TB vaccine construct model highlighted its structural integrity with more than 91.1% amino acid residues in the most favourable regions. Molecular docking of bovine TB vaccine construct and TLRs (TLR-2,4 and 6) was then undertaken. The results of docking analysis suggested stable and robust interactions between the TB vaccine construct and TLRs with binding energies of −55.15 kcal/mol (bovine TB vaccine-TLR2), −61.78 kcal/mol (bovine TB vaccine-TLR4) and −53.89 kcal/mol (bovine TB vaccine-TLR6). The strong interaction between the bovine TB vaccine construct and the host TLRs showed that the designed vaccine could initiate a robust immune response inside the host.

Next, subtractive proteomic analysis was performed to identify potential drug targets that could further help to investigate therapeutics drugs for the treatment of bovine TB. Some crucial steps were employed in the study to discover unique drug targets that are pivotal for the survival of *Mycobacterium bovis* and do not initiate hypersensitive reactions or side effects in the host. The 1163 conserved proteins of *Mycobacterium bovis* AF2122/97 were utilized for the subtractive proteomics approach to identify drug targets. First, the homologous protein sequences between *Mycobacterium bovis* and the host (cattle) were excluded from the study. Thus, 735 non-homologous proteins remained out of the 1163 proteins. The subsequent analysis showed that more than 50% of the non-homologous proteins, specifically a total of 386 proteins, were involved in distinctive pathways in *Mycobacterium bovis.* Around 56.74% of these proteins were enzymatic, and 43.26% were non-enzymatic proteins.

An assessment of other crucial criteria, such as essentiality of a target for survival, virulence, and translation efficiency of the selected protein, to identify potential drug targets was then carried out. Out of 195 essential proteins predicted by DEG, 29 proteins were commonly predicted as virulent by the VirulentPred and MP3 tools. Of these 29, only 13 proteins were found to have a CAI score greater than 0.7. These 13 proteins selected were further analysed for their physiochemical properties, structural homology and druggability properties. Analysis of the physiochemical properties showed that all 13 proteins had characteristics of potential drug targets. Structural homology analysis showed that out of the 13 proteins, four proteins had no suitable template structure available in the PDB database and were excluded from the study. Finally, the druggability potential of the shortlisted nine drug targets was predicted using DrugBank. The six proteins (YP_009357346.1, YP_009357464.1, YP_009357822.1, YP_009358342.1, YP_009359528.1 and YP_009361152.1) with considerable similarity found in DrugBank were termed druggable target proteins. The remaining three (YP_009357462.1, YP_009358851.1 and YP_009359962.1) proteins with no significant hits found were considered novel target proteins. The nine highly potential drug targets identified in the study could facilitate the development of novel drugs against *Mycobacterium bovis*.

Thus, using an extensive and systematic approach, this study presented the development of an epitope-based vaccine that would be far more effective than the current BCG vaccine for bovine TB. It also found, from a comprehensive investigation, nine potential drug targets that could be far more effective than the currently used drugs in treating bovine TB. Our research thus makes novel contributions to the vaccine and drug development for bovine tuberculosis. Further, laboratory studies are suggested to validate the predicted vaccine and drug targets.

## Supporting information

https://lincolnuniac-my.sharepoint.com/:u:/r/personal/sandhya_samarasinghe_lincoln_ac_nz/Documents/Research%20R%20Drive/Sandhya%20Samarasinghe/D/(D)%2

